# TLK1-mediated RAD54 phosphorylation spatio-temporally regulates Homologous Recombination Repair

**DOI:** 10.1101/2022.09.19.508551

**Authors:** Ishita Ghosh, Youngho Kwon, Aida Badamchi Shabestari, Rupesh Chikhale, Jing Chen, Claudia Wiese, Patrick Sung, Arrigo De Benedetti

## Abstract

Environmental agents like ionizing radiation (IR) and chemotherapeutic drugs can cause severe damage to the DNA, often in the form of double-strand breaks (DSBs). Remaining unrepaired, DSBs can lead to chromosomal rearrangements, and cell death. One major error-free pathway to repair DSBs is homologous recombination repair (HRR). Tousled-like kinase 1 (TLK1), a Ser/Thr kinase that regulates the DNA damage checkpoint, has been found to interact with RAD54, a central DNA translocase in HRR. To determine how TLK1 regulates RAD54, we inhibited or depleted TLK1 and tested how this impacts HRR in human cells using a I*Sce*-I-GR-DsRed fused reporter endonuclease. Our results show that TLK1 phosphorylates RAD54 at three threonines (T41, T59, and T700), two of which are located within its N-terminal domain (NTD) and one is located within its C-terminal domain (CTD). Phosphorylation at both T41 and T59 supports HRR and protects cells from DNA DSB damage. In contrast, phosphorylation of T700 leads to impaired HRR and engenders no protection to cells from cytotoxicity and rather results in repair delay. Further, our work enlightens the effect of RAD54-T700 (RAD54-CTD) phosphorylation by TLK1 in mammalian system and reveals a new site of interaction with RAD51.

## Introduction

DNA double-strand breaks (DSBs) can be caused by endogenous or exogenous agents and can be toxic in eukaryotic cells. If unrepaired, DSBs can lead to chromosomal rearrangements, cell transformation, or cell death(1,2). DSBs can be two-ended or - in the context of a replication fork - one-ended. While the former form of a DSB can be repaired by either NHEJ or HRR, the latter is resolved predominantly by HRR(3). In G2 phase cells when induced with DSB, inter-sister chromatid recombination happens post-replication(4).

Human Tousled like Kinases are Ser/Thr kinases, which show highest activity in S-phase. The mammalian genome encodes two TLK homologs which are known as TLK1 and TLK2(5). These proteins share 89% homology across their entire amino acid sequence and 94% similarity in their C-terminal kinase domain(6). *TLK1* gene has many splice variants (e.g TLK1B) that are expressed in a cell-specific context or induced in a stress-dependent manner, as identified from our previous work(7-9). TLK1B lacks the first 237 N-terminal amino acids but the kinase domain is conserved. Previous studies have shown that overexpression of either the full-length (TLK1) or the spliced variant (TLK1B) in mouse mammary fibroblast cells confers radio-resistance(10). Depletion of TLK1 delays S-phase progression, whereas dominant expression of a kinase-dead TLK1 leads to impaired DSB repair efficiency in cells during recovery from irradiation (IR), and pharmacologic inhibition of TLK with certain phenothiazines impairs DSB repair and leads to accumulation of γH2AX (11).

There are multiple identified substrates of TLK1 that serve in the DNA damage response pathway(7). TLK1 regulates DNA damage checkpoints (intra-S phase) through Rad9 phosphorylation at S328 which leads to dissociation of the 9-1-1 alternative clamp loader (12-14). TLK1 phosphorylates NEK1 at T141 and regulates its activity which further activates and stabilizes ATR-ATRIP complex(7). However, whether TLK1 plays a direct role in DSB repair is unknown.

In human cells, HRR depends on RAD54, an ATP-dependent DNA motor translocase(15). Phylogenetically, RAD54 belongs to SWI/SNF superfamily of helicases without canonical helicase function(16). The ATPase activity is required for double strand DNA translocation(17). After DNA damage during S/G2 phase, RAD54 interacts with the RAD51 recombinase and mediates HRR at different stages of the pathway (18,19). During the pre-synaptic stage, RAD54 stabilizes the RAD51 nucleoprotein filament (invading strand)(20) and facilitates homology searching (17). After establishing successful homologous contacts, RAD54 converts the synaptic complex into heteroduplex D-loop structures(21-23). At a later stage of HRR, RAD54 can facilitate branch migration of the Holliday Junction structures in an ATP-dependent manner, which ultimately are resolved to recombinant products through intricate processes(24). Additionally, RAD54 can disassemble RAD51 from heteroduplex DNA to allow DNA polymerase recruitment for repair synthesis(25,26). The unstructured (1-90 aa) and structured (91-154 aa) regions of the RAD54 N-terminal domain (NTD) physically interact with RAD51 *in vitro*(27). This interaction is conserved between yeast and human (15,23,28). The structured C-terminal domain (CTD) contacts the dsDNA backbone(29). Till date, there has been no evidence of the RAD54-CTD interacting with RAD51 *in vitro* or *in vivo*.

In this study we explore the role of TLK1 in HRR. We find that TLK1 interacts with human RAD54 and phosphorylates it at three novel residues, two within the NTD (T41 and T59) and one within the CTD (T700). We further find that phosphomimic RAD54-T41, T59D (RAD54-T2D) confers protection of cells from the cytotoxic effects of ionizing radiation (IR) and shows higher HRR capacity than its counterpart, while phosphomimic RAD54-T700 (RAD54-T700D) renders cells more radiosensitive and impaired in HRR. We report that the human RAD54-T700D mutant binds more tightly to RAD51 *in vitro* and *in vivo*. Moreover, homology molecular modelling indicates that Lys70 of RAD51 interacts with pT700 of RAD54. Our study reveals that human TLK1 phosphorylates RAD54 which can interact with RAD51 through a novel interacting CTD surface and thus negatively regulate HRR completion by deferring RAD51 disassembly.

### Materials and Methods Cell culture

HeLa cells were cultured in high glucose DMEM (Sigma Aldrich) with 10%FBS (Atlanta Biologicals) and 1x Antibiotic-Antimycotic (Gibco) and incubated in5% CO_2_ at 37°C in a humidified incubator. U2OS-DRGFP was a kind gift from Xiao-Fan Wang(30). All cells were cultured under same condition.

### DNA damage treatments

Cells were irradiated using X-Ray irradiator (Elekta Versa HD, linear accelerator) with a dose rate 400MU per min. Mitomycin C (Sigma Aldrich, cat# M0503), was solubilized in Phosphate Buffer Saline (PBS) as a stock concentration of 0.2mg/ml (0.6mM) and diluted to indicated concentrations in DMEM freshly prior to each experiment.

### Cell synchronization

Cells were seeded in 6 well plates at a density of 0.2X10^6 cells/well for cell-cycle analysis. 24hrs later cells were treated with 4mM HU for 16hrs. Media was replaced with fresh DMEM and cells were treated with IR (10Gy) and allowed to recover for indicated times. After each time-point either cells were collected for fractionation and immunoblotting or cell-cycle analysis. For cell-cycle profile, cells were washed with ice-cold PBS and fixed in 70% ethanol overnight at 4°C. Following day ethanol was removed and cells washed with fresh PBS and stained with Propidium Iodide (50 μg/ml) (Sigma Aldrich P4170) in a solution of 3.8mM Sodium Citrate and 0.5 µg/ml RNase A. Samples were analyzed in Flow-cytometer (Becton Dickinson) and quantitative amounts of cell cycle phase (G1, S, G2, G0) determined by FACS Diva Modfit software.

### Expression of recombinant protein and purification

Recombinant 6X-His-human TLK1 kinase domain (TLK1B) encoding plasmid was expressed in *E. coli* [Rosetta Gami (DE3)] and purified as previously published (31). TLK1 kinase activity was assayed using the ADP hunter kit as published (32). RAD54 was expressed in the form of His6-Thioredoxin-RAD54-Flag in Rosetta cells (Novagen) transformed with pET32-RAD54-Flag. The proteins were purified as described (33) following the purification protocol of *S. cerevisiae* Rad54 homolog (Rdh54).

### Identification of RAD54 phosphorylation by liquid chromatography-electrospray ionization-tandem mass spectrometry (LC-ESI-MS/MS)

All mass spectra reported in this study were acquired by the University of Kentucky Proteomics Core Facility. The recombinant purified proteins of RAD54 from in vitro kinase assay (with or without TLK1B) was run in 8% SDS-PAGE gel and proteins identified with Coomassie staining. The protein gel bands corresponding to RAD54 were excised and subjected to dithiothreitol reduction, iodoacetamide alkylation, and in-gel trypsin digestion using a standard protocol. The resulting tryptic peptides were extracted, concentrated and subjected to shot-gun proteomics analysis as previously described (34). LC-MS/MS analysis was performed using an LTQ-Orbitrap mass spectrometer (Thermo Fisher Scientific, Waltham, MA) coupled with an Eksigent Nanoflex cHiPLC™ system (Eksigent, Dublin, CA) through a nano-electrospray ionization source. The peptide samples were separated with a reversed phase cHiPLC column (75 μm x 150 mm) at a flow rate of 300 nL/min. Mobile phase A was water with 0.1% (v/v) formic acid while B was acetonitrile with 0.1% (v/v) formic acid. A 50 min gradient condition was applied: initial 3% mobile phase B was increased linearly to 40% in 24 min and further to 85% and 95% for 5 min each before it was decreased to 3% and re-equilibrated. The mass analysis method consisted of one segment with 11 scan events. The 1st scan event was an Orbitrap MS scan (300-1800 m/z) with 60,000 resolution for parent ions followed by data dependent MS/MS for fragmentation of the 10 most intense multiple charged ions with collision induced dissociation (CID) method.

### MS/MS protein Identification

The LC-MS/MS data were submitted to a local mascot server for MS/MS protein identification via Proteome Discoverer (version 1.3, Thermo Fisher Scientific, Waltham, MA) against a custom database containing RAD54 (Q92698 [RAD54_HUMAN], used for RAD54 samples) downloaded from Uniprot. Typical parameters used in the MASCOT MS/MS ion search were: trypsin digestion with a maximum of two miscleavages, cysteine carbamidomethylation, methionine oxidation, as well as serine, threonine, and tyrosine phosphorylation. A maximum of 10 ppm MS error tolerance, and a maximum of 0.8 Da MS/MS error tolerance. A decoy database was built and searched. Filter settings that determine false discovery rates (FDR) are used to distribute the confidence indicators for the peptide matches. Peptide matches that pass the filter associated with the FDR rate of 1% and 5% are assigned as high and medium confident peptides, respectively.

### Site-directed mutagenesis (SDM)

HA tagged-RAD54-WT encoded plasmid (pcDNA3.1/neo) was used as template to create SDM using Agilent QuikChange Lightning Multi Site-Directed Mutagenesis Kit (Cat #210515). PCR conditions were used as manufacturer’s protocol with following SDM primers.

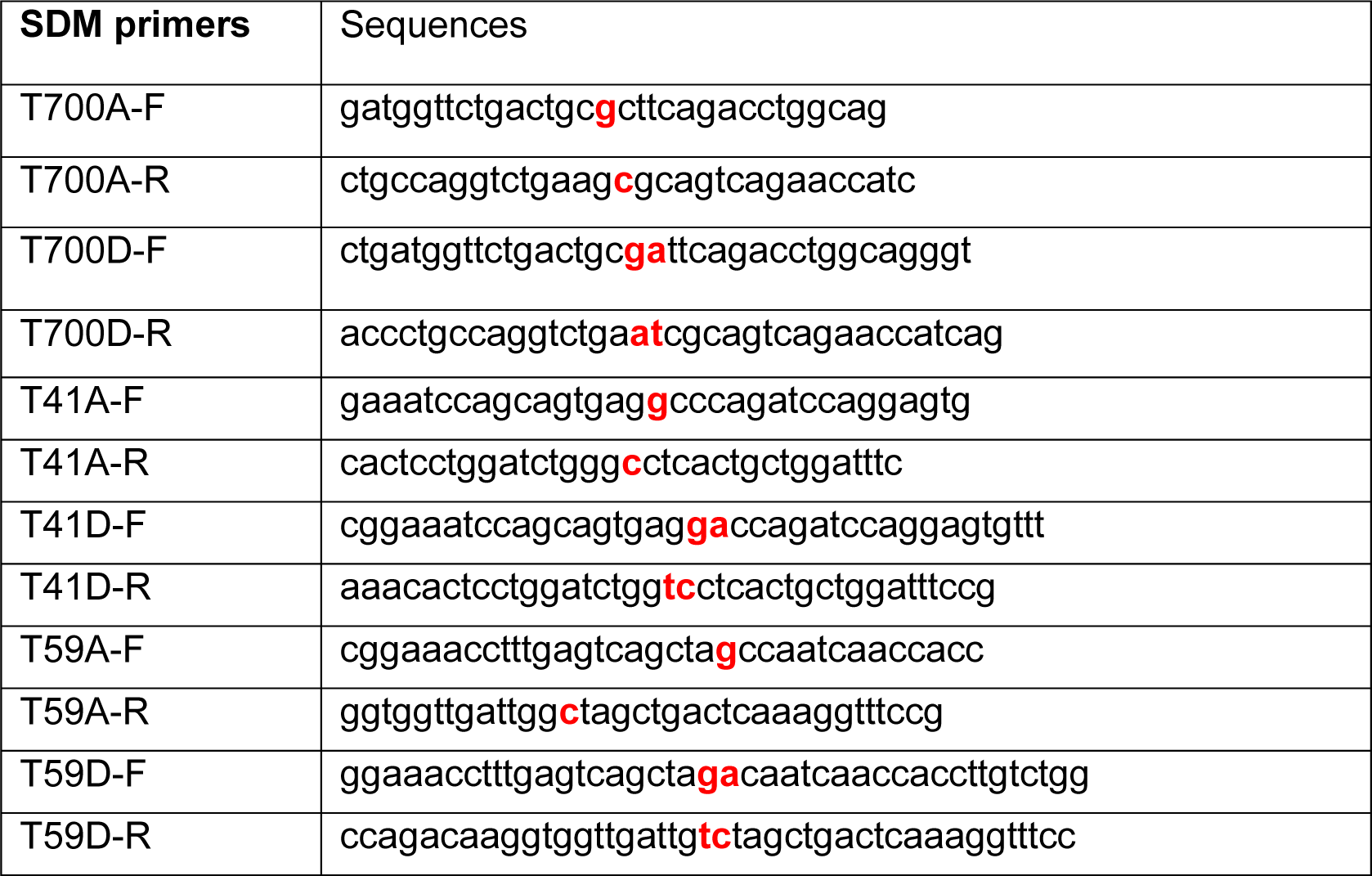

### Generating stable cell lines

HeLa *RAD54KO*-DRGFP was generated by transfecting DRGFP plasmid. pDRGFP was a gift from Maria Jasin (Addgene plasmid # 26475; http://n2t.net/addgene:26475;RRID:Addgene_26475). A stable cell line was generated by selecting transfected cells with 2μg/ml Puromycin for three weeks. Media was replaced every three days with fresh media containing puromycin. HeLa *RAD54KO*-DRGFP cells were transfected with HA tagged-RAD54-WT encoded plasmid (pcDNA3.1/neo) or T700A/D or T2A/D mutants with a C-terminal HA-tag using Fugene HD (Promega). The RAD54-GFP encoding plasmid (pEGFP-Rad54-N1) was a kind gift from Ronald Kanaar (Erasmus University Rotterdam, Netherlands)(18). HeLa *RAD54KO* cells were transfected with 3μg of RAD54-GFP or eGFP plasmid per well in a 6-well plate using Lipofectamine LTX PLUS (Invitrogen). Stable cell lines were generated by selecting with neomycin (800μg/ml) for three-four weeks. HeLa *RAD54KO-*RAD54-GFP or HeLa *RAD54KO-*eGFP cells were sorted by flow-cytometry and GFP+ve cells were seeded to generate the clones.

### DRGFP assay

I-SceI-GR-DsRed was a gift from Tom Misteli (Addgene plasmid # 17654; http://n2t.net/addgene:17654;RRID:Addgene_17654). Transient transfection was done using Fugene HD for 24hrs according to manufacturers protocol (Promega Corp). 3μg of I-Sce-GR-RFP plasmid DNA was diluted in OptiMEM. Fugene was added per well in a ratio [DNA (μg): Fugene (μl)= 1:3]. 150μl of Fugene+DNA+OptiMEM mixture was added to each well and cells were incubated for 15mins. Later, 1.5ml of DMEM (without antibiotic) was added to cells and incubated for 24hrs. After 24hrs, Triamcinolone acetonide (TA) was added to cells and incubated for 20mins at a final concentration of 0.1μM in DMEM (without antibiotic) to induce SceI localization to nucleus. After 20mins, media was replaced with complete DMEM and cells were incubated for 48hrs. After 48hrs, RFP+ve cells were counted in Flow-cytometer and GFP+ve cells out of RFP+ve population of cells measured. Control (No SceI) transfected cells were used for RFP and GFP gating.

*TLK1 depletion for DRGFP assay*-TLK1 depletion from HeLa or HeLa-DRGFP cells were performed using 250nM shRNA of TLK1. TLK1 shRNA (ATTACTTCATCTGCTTGGTAGAGGTGGCT) plasmid was obtained from origene (Rockville, MD, USA, cat# TR320623). Transfection was done using Fugene as per manufacturer’s protocol for plasmid DNA. Stable lines were generated by selection with Puromycin for two weeks. DRGFP assay was performed as mentioned before.

*TLK1 inhibition* was done using 10uM J54 (iTLK1) in U2OS-DRGFP cells 24hr post I-SceI induction with TA. Flow-cytometry was done 24hrs later.

For all the HeLa-RAD54KO expressing RAD54 mutants, I-SceI transfection was performed as mentioned before.

*TLK1 and 53BP1 depletion by siRNA –* 0.3X10^6 U2OS-DRGFP cells were seeded in 6-well plate 24hrs prior to siRNA transfection. si53BP1(5’-GAAGGACGGAGUACUAAUA-3’) and siTLK1 (5’-GAAGCUCGGUCUAUUGUAA-3’) purchased from Sigma-Aldrich and transfected for 24hrs in cells using RNAimax as per manufacturers protocol (Invitrogen). I-SceI-GR-DsRed transfection was done the following day as mentioned above.

### Immunoprecipitation assay

For human TLK1 and RAD54 or RAD51 protein interaction, Protein A/G PLUS agarose (cat# sc-2003, Santa Cruz Biotechnology) beads (100μl slurry or 50μl packed beads per reaction) equilibrated in 1X PBS were used to coat with anti-TLK1 or IgG isotype antibody (0.5μg antibody per reaction, cat# 720397, Thermofisher). To ensure uniform coating of beads, the beads were incubated with TLK1 antibody overnight in rotor at 4°C in presence of 500μl PBS. On the following day, TLK1 +beads were centrifuged at 1000g for 1min at 4°C and the supernatant was discarded. BSA (0.5μg) was added to each reaction to increase specific interaction between TLK1 protein and its cognate antibody. Reactions were incubated for 30mins in a tumbler at 4°C. Equal amounts (0.5μg) of TLK1 (2.5μl of TLK1B concentration= 0.25μg/μl) and RAD54 (5μl of rc-RAD54= 0.09μg/μl) or human RAD51 (1μl of rc-RAD51= 0.5μg/μl) proteins were loaded for IP reactions and volume made up to 200μl with PBS +0.1% Tween20 (PBST). The reactions were incubated in a tumbler at 4°C for 2hrs. After 2hrs, samples were centrifuged at 1000g for 1min followed by aliquoting the supernatant (loaded as S fraction) and beads were washed three times with PBST. Proteins were eluted with 2X SDS-Laemmli buffer (25μl) and boiled for 5-6mins. After a short-spin, 25μl samples were loaded per lane and analyzed by western blotting (loaded as E fraction). For input samples, the same amount of proteins were analyzed in parallel For RAD54 co-immunoprecipitation from HeLa cell lysates, cells were lysed with RIPA buffer and sonicated at 10secs ON/OFF pulse for 3cycles. For IP reactions, RAD54 antibody (cat# sc-166730, Santa Cruz Biotechnology) or IgG (mouse) isotype (2μg of antibody) was incubated with pre-equilibrated Protein A/G agarose beads (cat# sc-2003, Santa Cruz Biotechnology) in RIPA buffer for 4hrs at 4°C in rotor. One mg of total protein was loaded for each reaction and volume made up to 1ml with RIPA and incubated overnight at 4°C in a rotor. Following day, samples were centrifuged at 1000g for 1min and then washed thrice with RIPA. Eluted with 25μl 2X SDS-laemli buffer. Samples were boiled for 5mins and loaded in 8% SDS-PAGE gel for immunoblot analysis.

For RAD54-HA immunoprecipitation, equilibration of Pierce anti-HA agarose bead (cat#26181, Thermo Scientific) (∼30μL packed volume per reaction) were washed with Pierce IP lysis buffer (cat# 87787, Thermo Scientific) [25 mM Tris-HCl pH 7.4, 150 mM NaCl, 1% NP-40, 1 mM EDTA, 5% glycerol] three times. The resuspended beads were centrifuged at 1000rpm for 30secs at 25°C. The beads were then resuspended in 60μL of the buffer and cell lysate (1.0mg of total proteins) and protease/phosphatase inhibitor cocktail (final 1X) was added in 1ml IP lysis buffer. Samples were incubated in rotor for 2hrs at 4°C. After 2hrs, tubes were centrifuged for 30secs at 1000rpm at 25°C. The supernatant was discarded and the beads were washed with 500μL of IP lysis buffer three times. Samples were eluted using 30μL of 2X SDS laemmli buffer to each tube and boiled for 5mins. Same volume was loaded in 8%SDS-PAGE gel for immunoblot analysis. For input samples, 50μg of total proteins were diluted in 4X SDS-laemmli buffer and loaded in same gel.

### Clonogenic assay

HeLa RAD54KO, HeLa RAD54KO WT (reconstituted), HeLa RAD54KO-T700A/D, HeLa RAD54KO-T2A/D cells in suspension were treated with indicated IR doses and 250, 500 and 1000 cells were seeded in triplicate. Cells were allowed to grow for 10days for colony formation. Cells were fixed with Crystal violet solution (3gm in 5% Methanol: 5% Isopropanol) (BD BBL Gram Crystal Violet, Cat # 212525) and counted manually using ImageJ. For MMC clonogenic assay, 250, 500 cells were seeded in each well in triplicate. 24hrs later cells were treated with indicated doses of MMC. 6hrs later media was replaced with fresh DMEM and cells allowed to grow for 10days followed by crystal violet staining.

### pRAD54-T700 custom antibody generation

p-RAD54-T700 site-specific antibody was generated by Thermofisher Scientific, Life Technologies designed against phosphorylated Thr residue (bold) in peptide (PDGSDC**T**SDLAGW).

**ATPase Assay**-The RAD54 protein (50nM) was incubated in 20μl of the reaction buffer (35 mM Tris-HCl, pH 7.4, 22.5 or 75mM KCl, 1mM dithiothreitol, 5mM MgCl_2_, 100ng/μl BSA, 5ng/μl pBluescript dsDNA, and 0.4μM ATP supplemented with 5nCi/μl [gamma-32P]ATP) at 37°C for the indicated times. To examine the effect of RAD51, 200nM RAD51 was added in the reaction. The reactions were stopped with an equal volume of 500mM EDTA. The level of ATP hydrolysis was determined by thin layer chromatography, followed by phosphorimaging analysis using Typhoon PhosphorImager and quantified with Image Quant (Cytiva).

### Western Blotting

Total 40μg of protein from cell lysates were loaded in 8%SDS-PAGE gel and immunoblotted with RAD51 antibody (1:1000; PA5-27195, Thermofisher Scientific) diluted in 1%BSA+TBS+ 0.1%Tween 20, RAD54 antibody (1:1000; sc-374598; sc166370, Santa Cruz Biotechnology) diluted in TBS+0.1%Tween 20, HA tag antibody (1:1200; cat# 04-902 clone DW2, EMD Millipore), GFP antibody (1:2000; cat# MA5-15256 (GF28R), Thermofisher Scientific), 53BP1 antibody(1:1000; cat# MAB3804 clone BP18, EMD-Millipore), Ku 70 (1:1000, cat# sc-17789 (E-5), Santa Cruz Biotechnology), ORC2 (1:1000, cat# PA5-67313, Thermofisher Scientific). Blots incubated overnight at 4°C. HRP-conjugated goat anti-rabbit or goat anti-mouse IgG (1:2000; Cell Signaling Technology) were used as secondary antibodies and blots incubated for 1hr at room temperature. Blots were developed using ECl chemi-luminescence substrate and imaged in Chemidoc (Touch) system v2.3.0.07, Bio-Rad. Western blot signals were quantified using Image Lab software v6.0.1 (Bio-Rad).

### Cellular fractionation

10^7 HeLa cells were seeded 24hrs prior to treatment. After treatment, cells were harvested. Nuclear fractions were isolated in a method published previously(13). Briefly, ∼10^7 cells were resuspended in buffer A (10mM HEPES [pH 7.8], 10mM KCl, 1.5mM MgCl2, 0.3M Sucrose, 10% Glycerol and 1mM Dithiothreitol (DTT)) supplemented with halt EDTA free protease and phosphatase inhibitors (Life technologies, Cat# 78441). 0.1% Triton X-100 was added and cells incubated for 5mins on ice. Cytosolic proteins were separated from nuclei by centrifugation at 1300xg, for 4mins. Nuclei were washed once in buffer A and then lysed in buffer B (20mM Tris [pH 8.0], 3mM EDTA, 0.2mM EGTA and 1mM DTT) supplemented with halt EDTA free protease and phosphatase inhibitors for 30mins. Insoluble chromatin was then separated from soluble nuclear proteins by centrifugation at 1700xg for 4mins, washed once in buffer B and repeated centrifugation. The final chromatin pellet was resuspended in 1X RIPA buffer to release chromatin bound proteins and incubated on ice for 5mins followed by bio-ruptor sonication at high amplitude for 10cycles. Protein sample collected by centrifugation at 10000g for 5mins. The fractions were analyzed by 8%SDS-PAGE followed by western blot with Ku-70 and ORC2 antibodies.

### Proximity Ligation Assay (PLA)

Cells were seeded at 0.1X 10^6 cells/ well on 10mm coverslips coated with poly-Lys in a 24-well plate 24hrs before treatment with 10Gy IR. After treatment, cells were washed twice with 1X PBS at room temperature. Cells were fixed with 4% PFA for 15mins at room temperature in dark. Cells were then washed twice with PBS and permeabilized with PBS+ 0.2% Triton X-100 for 10mins. Cells were washed once with PBS and then blocked with 1X Duolink blocking solution (Cat # DUO92101, Sigma-Aldrich) for 1hr at 37°C. After blocking buffer was removed, primary antibody (anti-RAD54; sc-374598 and anti-RAD51; ab-176458, Abcam) was diluted in PBS +0.1%(v/v) Tween 20 +2% (w/v) Bovine Serum Albumin (BSA) and incubated overnight at 4°C. The next day, primary antibody removed, and cells washed twice with 1X wash buffer A [0.01M Tris-HCl (pH 7.4), 0.15M NaCl and 0.05% Tween 20], 5mins each. Cells were incubated in a humidified chamber with Duolink in situ PLA PLUS and MINUS probes (Sigma Cat # DUO92101) diluted 1:5 in PLA antibody diluent. Samples were washed twice with 1X wash buffer A and ligase added (diluted 1:40 in ligation buffer). Incubated for 30mins in humidified chamber at 37°C. Amplification buffer was diluted from 5X to 1X. After timepoint, samples were washed twice with wash buffer A. Polymerase (Cat # DUO92101, Sigma-Aldrich) added at a dilution of 1:80 in 1X amplification buffer. Samples were incubated in humidifier for 100mins at 37°C. Polymerase was removed, and samples washed with 1X wash buffer B [0.2M Tris-HCl (pH 7.4), 0.1M NaCl) twice for 10mins each. Next, samples were incubated at 0.01X wash buffer B for 1min. Coverslips mounted using Duolink PLA mounting media with DAPI and incubated for 15mins before imaging. Images were acquired in Zeiss Axioobserver with 40X, oil objective.

### Indirect Immunofluorescence

Cells were seeded at 0.3X 10^6 cells/ well on 22mm coverslips coated with poly-L-Lys in a 6-well plate 24hrs prior to treatment. After treatment with irradiation or MMC as indicated, media was removed and cells washed with 1X PBS twice.

For RAD54 and RAD51 foci experiments, ice-cold cytoskeleton (CSK) buffer [(10mM Pipes, pH 7.0, 100mM NaCl, 300mM sucrose, and 3mM MgCl2) containing 0.5% Triton X-100] was added to cells for 4-5mins. After one PBS wash, cells were fixed with 4%PFA in PBS (cold) was added at room temperature (RT) for 15mins in the dark. PBS wash done twice. Cells were permeabilized with PBS +0.1% Triton X-100 for 10mins. PBS wash done twice 1-2mins each. Blocking was done with PBS +0.1%(v/v) Tween 20 +2% (w/v) BSA for 1hr at RT. Primary antibody for RAD54 (sc-374598, 1:250) and RAD51 (ab-176458, 1:500) were diluted in PBS +0.1%(v/v) Tween 20 +2% (w/v) BSA and incubated overnight at 4°C. Next day, primary antibodies were removed, and samples washed twice with PBS (1min each). Secondary antibodies [Goat anti-Mouse IgG (H+L) Alexa Fluor™ 488, Invitrogen (cat# A11029) and Goat anti-Rabbit IgG (H+L) Alexa Fluor™ 594, Invitrogen (cat# A11012)] were diluted at 1:1000 in PBS +0.1%(v/v) Tween 20 +0.1% BSA and added to samples for 1hr at room temperature. After 1hr, antibodies removed, and samples washed with PBS thrice. Coverslips mounted on pre-cleaned slides with anti-fade mounting media containing DAPI (vector labs, vectashield).

For TLK1 and γH2A.X or 53BP1 foci studies, 0.3X 10^6 cells/ well were seeded on 22mm coverslips coated with poly-L-Lys in a 6-well plate 24hrs prior to treatment. Cells were treated with MMC (3μM) for 2hrs and allowed to recover for 4hrs after which media was removed. Cells were washed in PBS twice. PBS+0.1% Triton X-100 (cold) added for 1min followed by one PBS wash. Cells were fixed with 4%PFA in PBS (cold) at room temperature for 15mins in dark. After two PBS wash, cells permeabilized with 0.5% Triton X-100 for 5mins at RT. PBS wash twice. Blocking was done with PBS +0.1%(v/v) Tween 20 +2% (w/v) BSA for 1hr at RT. PBS washed twice. Cells incubated with primary antibodies, anti-TLK1(cat# 720397, Invitrogen) and anti-γH2A.X (pS139) (cat# 05-636, EMD Millipore) or 53BP1 (1:500, cat# MAB3804) diluted in PBS+0.1%(v/v) Tween 20 +2% (w/v) BSA at ratio of 1:250 and incubated overnight at 4°C. PBS wash done twice. Secondary antibodies diluted as before and incubated for 1hr at RT. The DAPI staining and mounting done as mentioned previously.

RAD54-GFP analysis: HeLaRAD54KO reconstituted RAD54GFP cells were treated with iTLK1 (J54) and or IR and fixed at indicated timepoints. Cells were fixed using 4%PFA in PBS (cold) for 15mins at RT. Cells were PBS washed twice. Cells permeabilized using PBS +0.1%Triton X-100 for 10mins at RT. PBS washes done and stained with DAPI (1ug/ml) in PBSfor 5mins in RT. Cells mounted using Fluoroshield (Sigma, cat# F6182-20ML). Images acquired using Zeiss, Axiovision

### Image acquisition and analysis

Images were acquired in Zeiss Axioobserver, Axiovision, Apotome microscope equipped with Zen Blue 3.3 software (Carl Zeiss Microscopy). For PLA experiment imaging, 10 z-stacks with a step-size of 1μm optical distance with 12-bit per pixel, x-y dimension, 1388x1040 pixels were obtained. Image processing was done in Image J v2.3. Maximum intensity projection for each channel from all planes used for representation and quantification of foci. Quantification of foci was done using Andy’s algorithm(35) and nuclear foci and non-nuclear foci plotted with GraphPad Prism 9.

For all indirect immunofluorescence data, foci were quantified for each channel using image J after background correction. Foci were analyzed by Analyze particle tool and quantified after thresholding.

### Protein modelling and preparation

The RAD51 and RAD54L-WT proteins were obtained from the Alphafold database (https://alphafold.ebi.ac.uk/)(36), simultaneously these were also modelled for complete sequence of the RAD54L-WT and RAD54LT700D mutant using the MOE tool(37). The obtained models were compared structurally and then validated using the SWISSMODEL structure assessment for their structural quality based on Q-mean score, MolProbity check and Ramachandran analysis(38,39). These structures were later subjected to 500 ns molecular dynamics simulation to optimise the structural features(32). The optimised protein structures from these MD simulations were further subjected to structure assessment for their structural quality based on Q-mean score, MolProbity check and Ramachandran analysis.

### Protein-protein docking

The protein-protein docking was performed on the Haddock server (http://haddock.chem.uu.nl/) using standard protocol mentioned in the Haddock 2.2 docking manual(40). The protein files were uploaded on the server and parameters for number of structures to generate, protonation states for the histidine, definition of flexible segments, restraints, and associated parameters were defined. The randomisation of the starting orientation and protein minimisation parameters were also defined, mostly these were maintained at defaults. The results were obtained as clusters of top docking states and the top model was selected for further steps.

### Molecular Dynamics (MD) Simulation and Molecular Mechanics-Generalized Born Solvent Accessibility (MM-GBSA) Analysis

### System preparation

All the MD simulations were done on AMBER 18 software package(41,42). Protein complexes were prepared with the help of xleap. The RAD51-RAD54L-WT and RAD51-RAD54LT700D were solvated separately in truncated octahedron of TIP3P box(43). giving a total of 68787 and 63837 water molecules respectively. Sufficient number of counter ions Na+ and Cl-were added to neutralize the simulation system and 0.1M of ionic strength was achieved. To parameterize the amino acids and to model the proteins FF14SB force field was used(44).

### Unbiased MD simulation

Simulations were performed for each of the proteins RAD51, RAD54L-WT and RAD54L-Mutant, RAD54L-WT and RAD54LT700D complexes for 500 ns of time step on Nvidia V100-SXM2-16GB Graphic Processing Unit using the PMEMD.CUDA module. Simulations were run at 1 atm constant pressure using Monte Carlo barostat(45) and 300 K constant temperature by using Langevin thermostat with a collision frequency of 2ps-1 and the volume exchange was attempted for every 100 fs. An integration step of 2 fs was also used for simulation the hydrogen atoms involving bonds were constrained by using SHAKE algorithm(46). Long range electrostatic interactions were computed by using Particle Mesh Ewald method while for short range interaction a cutoff of 8 Å was used. Equilibration consisted of rounds of NVT and NPT equilibration for 10 ns in total. CPPTRAJ(47) was used to analyse the interactions over full trajectory after taking configuration at every 4 ps. RMSD, RMSF and MMGBSA binding free energy was determined after analysing the trajectories.

### Molecular Mechanics-Generalized Born Solvent Accessibility (MM-GBSA) analysis

The MM-GBSA(48) was performed on Amber18 and Amber18 tools. After simulation of the protein-protein complexes, another trajectory was obtained by continuing the MD simulation for 5 ns, this trajectory of 50 ns covering all the 2000 frames was used for MM-GBSA analysis. All the results in the form of energies were tabulated and reported in Kcal/mol.

## Results

### TLK1 regulates HRR and interacts with RAD54

We first wanted to study the effect of TLK1 on HRR. We transfected two different cell lines (HeLa and U2OS) containing the integrated DRGFP cassette with SceI-GR-DsRed plasmid. The transfection of SceI was measured by the red fluorescent signal and determines the transfection efficiency and expression of the nuclease whereas the green signal corresponds to the gene conversion event. TLK1 knockdown in HeLa-DRGFP cells by shRNA mediated silencing leads to a decrease of 50% HRR activity (Figure 1A, Figure S1A). A similar approach in U2OS-DRGFP cells with siRNA against TLK1 significantly shows 50% reduction in HRR whereas knockdown of 53BP1, an NHEJ promoting factor, shows an increase in HRR compared to respective control conditions (Figure S2A-C). In a complementary pharmacologic approach, inhibition of TLK1 in U2OS-DRGFP cells using a specific inhibitor, we find HRR activity decreases by 40% (Figure 1B, Figure S1B). These results indicate that TLK1 plays a role in HRR which remains to be explored. Next, we assessed the involvement of TLK1 in DSB repair by monitoring its recruitment to DSB repair foci marked by γH2A.X. We find that upon DSB induction, TLK1 foci show strong association with γH2A.X or 53BP1 foci, further suggesting that TLK1 plays an important role in DSB repair (Figure 1C-E and Figure S3).

**Figure 1:**
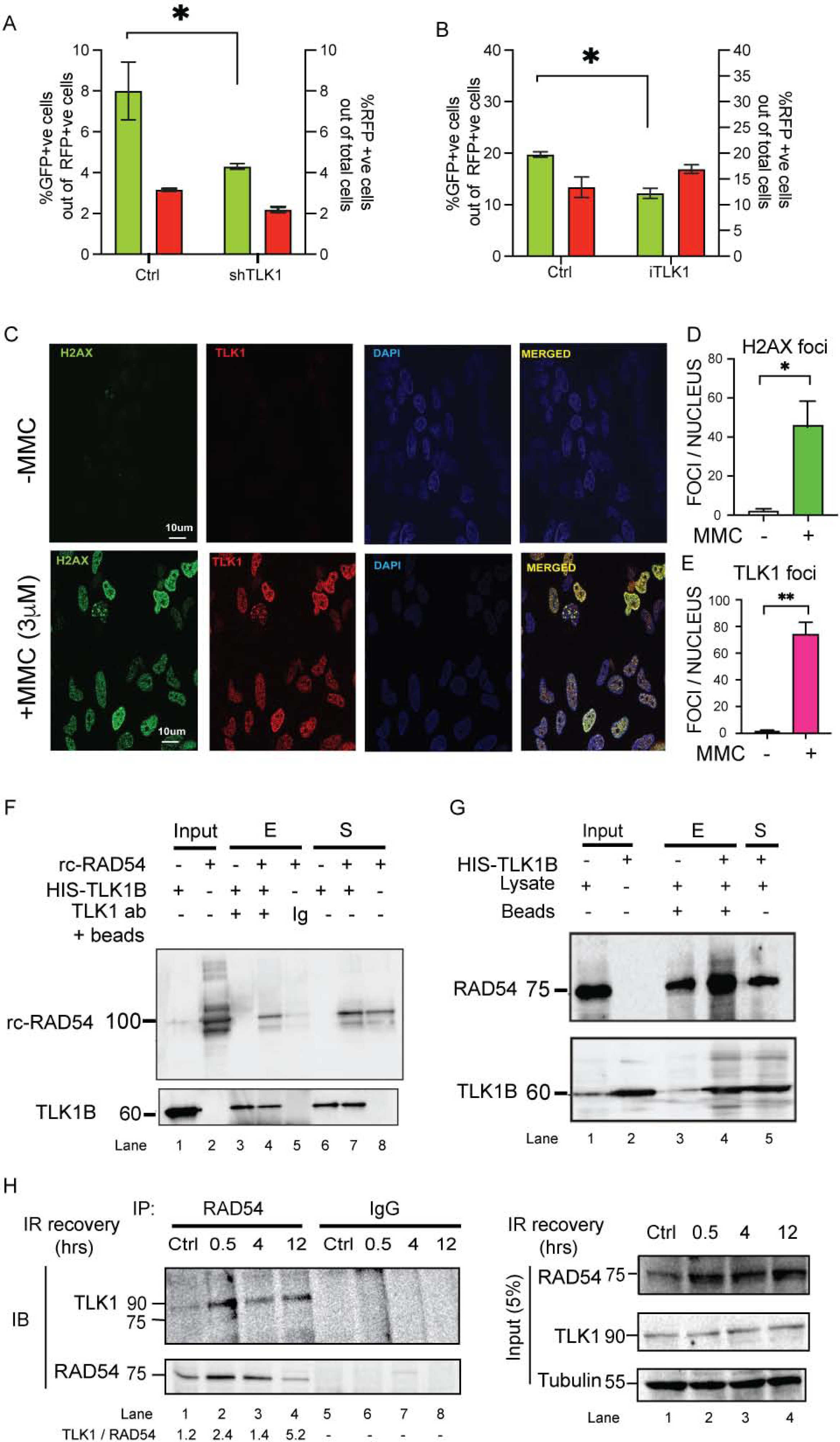
TLK1 regulates HRR and interacts with RAD54. A) TLK1 depletion in HeLa-DRGFP using shTLK1 (shRNA#1 shown) and B) TLK1 inhibition in U2OS-DRGFP cells (by iTLK1) lead to reduced in HRR efficiency. %GFP +ve cells (green bars corresponding to left y-axis) and %RFP+ve cells (red bars corresponding to right y-axis) as SceI transfected cell population in all cell lines (the results are mean ± SEM from 3 independent experiments. *, p<0.1; p-values were calculated from One-tailed Student’s t-tests (Unpaired) with Welch’s correction). C-E) TLK1 foci show strong correlation with γH2A.X (labelled-H2A.X) foci upon treatment with MMC (3μM). 100 nuclei from 3 independent experiments were assessed. Results are mean ± SEM. *, p<0.05, **, p<0.01; Unpaired Student’s t-tests with Welch’s correction. Scale bar is 10μm. F) TLK1B interacts with RAD54 (rc-RAD54); immunoprecipitation of recombinant TLK1B incubated with rcRAD54 protein using anti-TLK1- or IgG coated beads. G) HIS pulldown assay of HIS-TLK1B using Ni-NTA beads incubated with HeLa cell lysate. Interaction between HIS-TLK1B and RAD54 probed by immunoblotting. Upper panel, IB showing RAD54 enriched in TLK1B pulldown sample (lane 4, + HIS-TLK1B). Endogenous RAD54 (75kD) level in lysate (lane 3, - HIS-TLK1B). Lower panel, showing TLK1B band (60kD). Input lanes of lysate and TLK1B shown in lane 1 and 2. H) TLK1 interaction with endogenous RAD54 increases post-irradiation, as shown by co-immunoprecipitation (IP) reaction of RAD54 from HeLa cells treated with IR and allowed to recover for indicated times. TLK1 level normalized to RAD54 pulldown added in bottom panel. Right panel shows the input amounts for each reaction. Note, RAD54 expression is induced by DNA damage.

From our previous study(7), in which 9000 full-length human proteins were probed for interactors of TLK1, with recombinant human TLK1B, the top 10% interactors of the 160 proteins belonged to DNA damage repair pathways. Among these, we found that TLK1B interacts with RAD54B (RAD54 was not included in the protein array). RAD54B is a RAD54 paralog that shares significant homology (63%) with it(49,50). Therefore, we wanted to study if TLK1 can interact with RAD54. To this end, we used purified human RAD54 and the recombinant TLK1B splice variant in pull-down assays with TLK1 antibody and confirmed their association although some binding of RAD54 in the IgG isotype control was found probably because of non-specific RAD54 trapping in agarose beads (Figure 1F). We also tested for the presence of RAD54 in anti-TLK1 protein complexes generated in vitro with added His-TLK1B and HeLa cell lysates by co-immunoprecipitation (Figure 1G). Note that, we don’t find RAD51 interacting with TLK1B from recombinant proteins pulldown indicating that the interaction between TLK1 and RAD54 is not mediated by RAD51 (Figure S4). Next, we tested if the interaction between endogenous TLK1 and RAD54 is enhanced upon exposure of cells to DNA damage. We found that compared to the unirradiated control cells, cells treated with IR and subsequently allowed to recover for 0.5-12hrs post exposure show enhanced association of RAD54 with TLK1 (Figure 1H).

### TLK1 phosphorylates RAD54 at NTD and CTD to regulate HRR

To study the functional relationship between TLK1 and RAD54 and the possibility that TLK1 may engage in RAD54 phosphorylation, we performed *in vitro* kinase assays followed by LC-MS/MS. We obtained three novel TLK1-mediated phosphorylation sites in RAD54-two at NTD (T41 and T59) and one at CTD (T700), as shown in the schematic RAD54 domain map (Figure 2A). The phosphopeptides from m/z spectral peaks were analyzed against the control reaction (no TLK1) (Figure 2B and Figure S5, S6 and S7).

**Figure 2:**
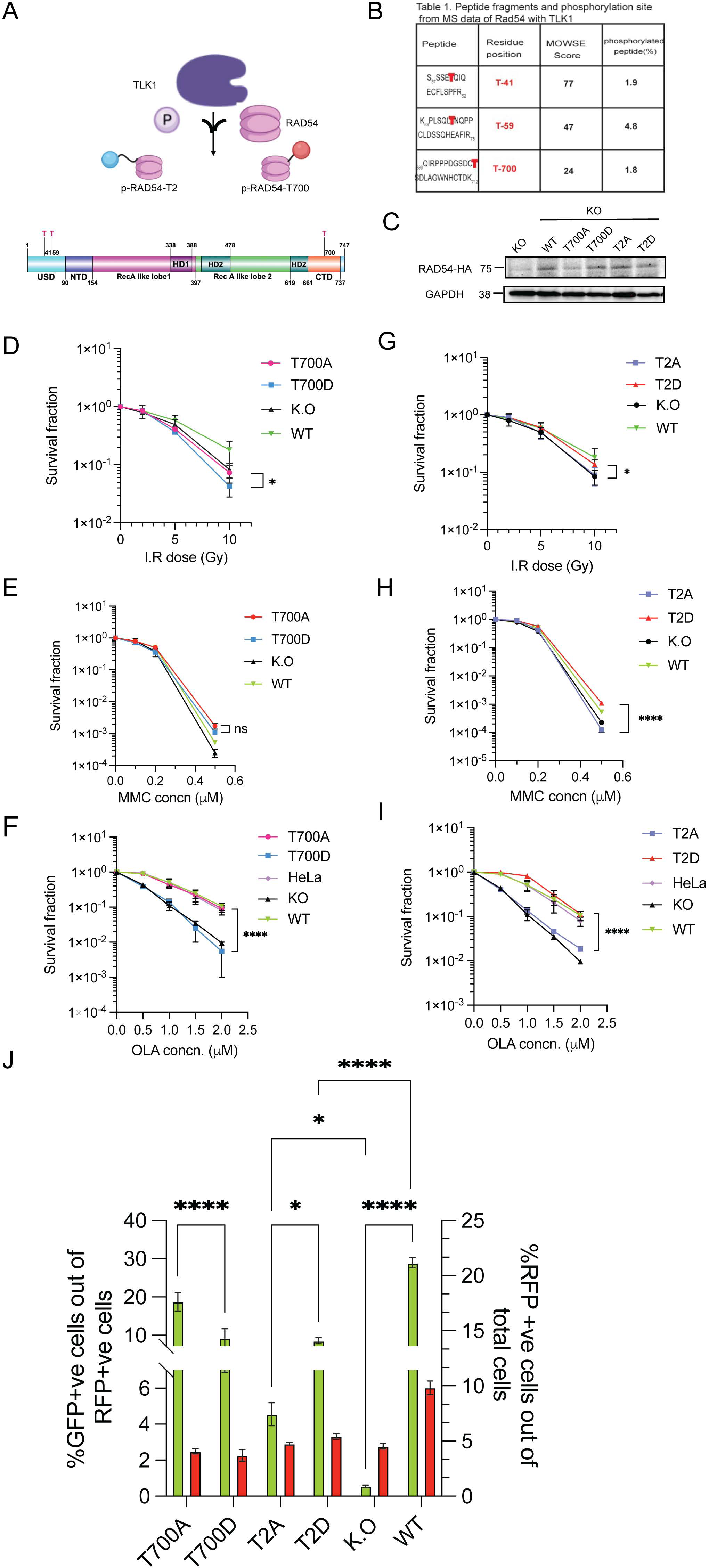
TLK1 phosphorylates RAD54 at NTD and CTD on novel sites and regulates HRR. A) TLK1 (purple lobed cartoon) phosphorylates (P) human RAD54 (in pink) at NTD (T41, T59 here shown together as T2, blue sphere) and CTD (T700; red sphere). Domain map of RAD54 showing sites of TLK1 phosphorylation on RAD54-NTD (T41 and T59) and RAD54-CTD (T700). B) LC-MS/MS data showing MOWSE scores on RAD54 site of phosphorylation. C) Expression of human RAD54 WT and phosphodefective (Ala-A) and phosphomimic (Asp-D), [at NTD (T2A/D) and CTD (T700A/D)] mutants in HeLa RAD54 K.O cells. D-F) Clonogenic assay of RAD54-T700A/D mutants after exposure to graded doses of IR, MMC and Olaparib (OLA) respectively and G-I) Clonogenic assay of RAD54-T2A/D post exposure to graded doses of IR, MMC and Olaparib (OLA) respectively. The results are mean ± SEM from three biological repeats *, p<0.05; ****, p<0.0001; Unpaired Student’s t-tests with Welch’s correction. J) DRGFP assay with RAD54 mutants in HeLaRAD54KO. Results showing comparison of %GFP +ve cells (left y-axis) in T700A, T700D, T2A, T2D with KO and WT reconstituted as controls and %RFP+ve cells (right y-axis) as I-SceI-GR-DsRed transfected cell population in all cell lines. Bars are the means ± SEM from three biological repeats. *, p<0.05; ****, p<0.0001; 2-Way ANOVA with Tukey’s multiple comparison was performed (simple effect within columns).

To dissect the function of TLK1-mediated RAD54 phosphorylation with a genetic approach, we changed the three threonines (Thr) to phospho-defective (Ala) or phospho-mimetic (Asp) residues using site-directed mutagenesis and generated stable cell lines in HeLaRAD54KO (51) cells expressing the ectopic RAD54 versions (Figure 2C and Figure S9A). Next, we asked if the RAD54-NTD mutants i.e RAD54-T41, 59A/D (RAD54-T2A/D) and the RAD54-CTD mutants (RAD54-T700A/D) mutants would show altered sensitivity to DSB-inducing agents. First, we performed clonogenic cell survival assays upon exposure of cells to IR. Our results show that, unlike the wild type protein, neither RAD54-T700A nor RAD54-T700D could rescue HeLaRAD54KO cells from the cytotoxic effects of IR (Figure 2D). In fact, expression of RAD54-T700D further sensitized HeLaRAD54KO cells to IR, suggestive of a dominant-negative effect. A similar trend was observed with MMC induced ICL damage for 6hrs indicating that the RAD54-T700 phosphomimic does not have a positive effect in ICL repair (Figure 2E), rather improved survival was observed between T700A and T700D (Figure S8A). Even the RAD54WT did not show significant difference in sensitivity from KO. The possible explanation could be the compensatory effect by paralog RAD54B on survival during MMC induction (52). Another consideration is that NHEJ is the primary mechanism of DSB repair in asynchronous HeLa cells. We chose a concentration range of 0.1-0.5μM MMC for clonogenic survival because at higher concentration of MMC (greater than 1μM), none of the lines survived. HR defect in cells selectively manifests sensitivity to PARP inhibitors. Previous studies have reported that RAD54KO has extreme sensitivity to Olaparib (52). A remarkable difference was observed between T700A and T700D in the treatment with Olaparib, where T700D had increased sensitivity slightly higher than RAD54KO (Figure 2F), while T700A showed significantly decreased sensitivity similar to WT (and parental HeLa) (Figure 2F). Clonogenic assays with the RAD54-T2A/D mutants showed that T2D mutant had higher survival capacity than T2A after IR, MMC or Olaparib induced damage that was comparable to that of WT reconstituted cells (Figure 2G, H and I, Figure S8B, S8C). These data suggests that the RAD54-NTD phosphorylation has an overall DNA DSB repair promoting function. At least, the RAD54-NTD mutants protects the human cancer cell line whereas the RAD54-CTD non-phosphorylated form has a similar function towards survival from Olaparib induced damage.

To better establish this, we tested the effects of these phosphomutants on HRR activity at DRGFP integrated into HeLaRad54KO cells after transfection with I-*Sce*I-GR-DsRed plasmids. Forty-eight hrs post SceI induction we counted GFP+ve cells in flow-cytometer. Our results show that T700A has 50% higher GFP+ve cells than T700D (Figure 2H) and similar qualitative expression was observed from fixed-cell microscopy Figure S8C). Although T700D has a 10% higher GFP conversion than RAD54KO cells and 50% lower than WT, T700D is not completely defective in HR whereas T700A has efficient HR activity with GFP conversion frequency similar to WT (20% GFP conversion). In a similar approach with T2A and T2D in the DRGFP assay, we find that T2D has 50% higher gene conversion frequencies than T2A from flowcytometry (Figure 2H) and also from microscopy (Figure S8D). While T2D has overall 10% GFP conversion, T2A shows only 4-5%GFP conversion. These results suggest that TLK1 mediated RAD54 phosphorylation at CTD perturbs HRR, while the NTD phosphorylation leads to efficient HRR in cells. Note that all the mutants and WT-complemented cells expressed very similar amounts of the ectopic RAD54 proteins (Figure 2C and Figure S9A).

### RAD54 phosphorylation at T700 increases post irradiation

Since the T700 site lies in the RAD54 CTD that is structurally known to interact with dsDNA donor template, we asked if this site is phosphorylated post IR damage in cells. In order to test the status of phosphorylation in cells, we generated an antibody against RAD54-T700 phosphopeptide. We tested the specificity of the antibody using the reconstituted WT, T700A and T700D cell lysates. The antibody generated specific signal against WT and T700D (phosphomimic mutants are often detected with P-specific antibodies(53)), but not T700A (Figure 3A). Next, we tested if the T700 phosphorylation is TLK1 specific. We treated the cells with TLK1 inhibitor, J54, which led to reduced signal intensities by ∼ 50% (Figure 3B, C). This is similar to the reduction observed for pNek1-T141 that is our standard marker to monitor TLK1 activity(32,54). Further, we asked if RAD54-T700 is phosphorylated post IR. To test this, we treated HeLa cells with IR (10Gy) and allowed cells to repair for 4hrs and 12hrs. We find that RAD54 is progressively phosphorylated at T700 as a function of time post IR (Figure 3D and Figure S10A). We obtained similar results in another cell line, U2OS (Figure S10B). These results suggest that RAD54 is phosphorylated by TLK1 at T700 primarily at a late stage of recovery from IR. We further wanted to test if RAD54 phosphorylated fraction is enriched in the nucleus of cells when cells are in S-phase post IR induction. We synchronized cells in S-phase (Cell cycle profile shown in Figure S11) and found that pRAD54 is increased at 4hrs and 12hrs post DNA damage (Figure 3E, lane 4 and 6, quantification in Figure 3F). When probed for RAD51, we find that RAD51 fractions are enriched in chromatin fraction at later IR recovery stage (12hrs) (Figure 3E, lane 6, quantification in Figure 3G). From all these observations we conclude that phosphorylated RAD54 and RAD51 association with chromatin increases at late DSB recovery stage.

**Figure 3:**
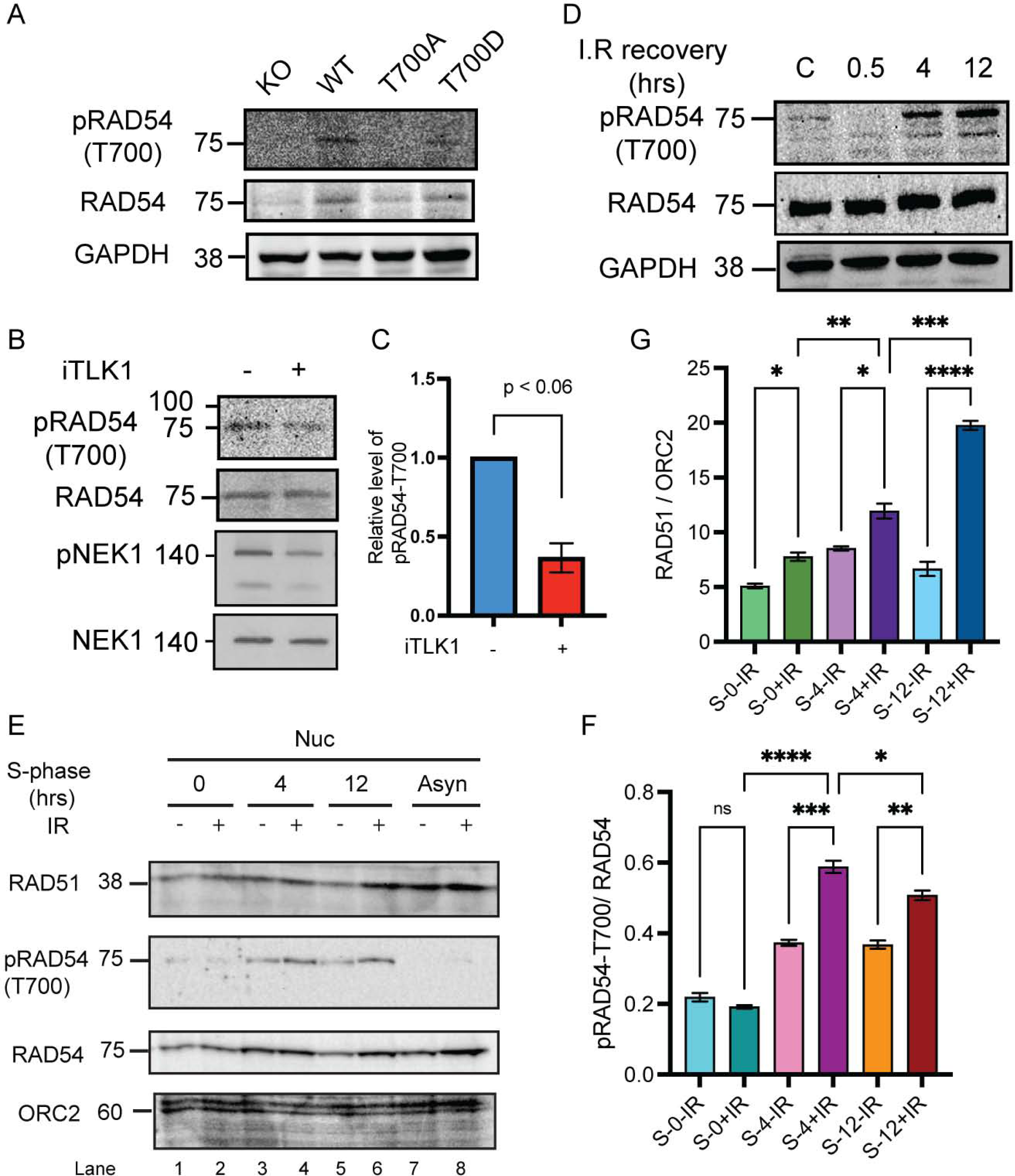
RAD54 phosphorylation occurs at T700 (pRAD54-T700) in cells in a TLK1 dependent manner and increases post IR recovery. A) pRAD54-T700 antibody specificity characterized to detect T700 site. B) pRAD54-T700 decreases by 60% after TLK1 inhibition. A parallel blot was probed for pNek1-T141 and Nek1. C) Quantification of relative level of pRAD54-T700 phosphorylation level from panel B. D) pRAD54-T700 is initially lost at 0.5hr, but increases post IR recovery (4 and 12hrs). E) Phosphorylated RAD54-T700 and RAD51 association increases in nucleus post IR induction in S-phase synchronized cells. RAD51 and p-RAD54-T700 probed in nuclear fraction of S-phase cells post DNA damage recovery as indicated. ORC2 is shown as chromatin associated nuclear loading control. Lane 1-2, 3-4, 5-6 indicates nuclear (Nuc) fractions from S-synchronized cells without or with IR (10Gy) and allowed recovery period of 0, 4, 12hrs respectively. Lane 7-8 indicates asynchronous (Asyn) cells. F) Quantification of pRAD54-T700 in S-phase nuclear fraction is shown. G) Quantification of RAD51 in S-synchronized nuclear fraction is shown. *, p<0.05; **, p<0.01; ***, p<0.001; ****, p<0.0001; Ordinary One-Way ANOVA followed by Tukey’s post-hoc analysis was performed for multiple group comparison.

### TLK1 phosphorylates RAD54 and alters RAD54 cellular localization

The localization of RAD54 in human cells under unperturbed condition has not been extensively studied. However, it is well known that RAD54 localizes to nucleus upon induced DNA damage(55,56). Here, we asked if RAD54 localization to nucleus is dependent on TLK1 phosphorylation. To do so, we first expressed RAD54-GFP in HeLaRad54KO cells (Figure S9B) and find that under control (non-irradiated) condition RAD54-GFP localizes to cytoplasm (Figure 4, control panel). After 2hrs recovery from 10Gy IR, with or without TLK1 inhibitor (J54, 10μM), we find that most RAD54-GFP shuttles to the nucleus (Figure 4, 2hrs IR recovery, -/+J54 panel). This shows that RAD54-GFP localization to nucleus following IR induction does not depend on TLK1 mediated RAD54 phosphorylation. However, after 10hrs of recovery when RAD54 returns to cytoplasm in control (Figure 4, 10hrs IR recovery, -J54 panel), TLK1 inhibition hinders the relocalization of RAD54-GFP to cytoplasm (Figure 4, 10hrs IR recovery, +J54 panel). Since there are 2 NTD and 1 CTD sites of TLK1-mediated phosphorylation that could govern the shuttling of RAD54, and we only have an antiserum for the pT700, we investigated the phosphorylation at this site in greater depth by cellular fractionation. In Figure S12A, we verified that total RAD54 was accumulating back in the cytoplasm only after 10hrs of IR recovery, and that J54 partly prevented this at 2hrs (Figure S12A, lane 9 vs lane 10). Remarkably, the pT700 antiserum revealed that the pT700-RAD54 is all nuclear, showing no signal in the cytoplasmic compartment. Whatever fraction of it is phosphorylated at T700, remains (or accumulates) in the nuclei after IR. And notably, J54 did not affect this in reducing its phosphor-signal, possibly best explained by the lack of a nuclear phosphatase associated with pRAD54 during the period of recovery from IR. Reprobing the blot with pNEK1 antiserum revealed that, in fact, J54 could reduce the signal from pNEK1-T141 (a specific activating phosphorylation by TLK1) even though this species was also primarily localized in the nuclear fraction (Figure S12B).

**Figure 4:**
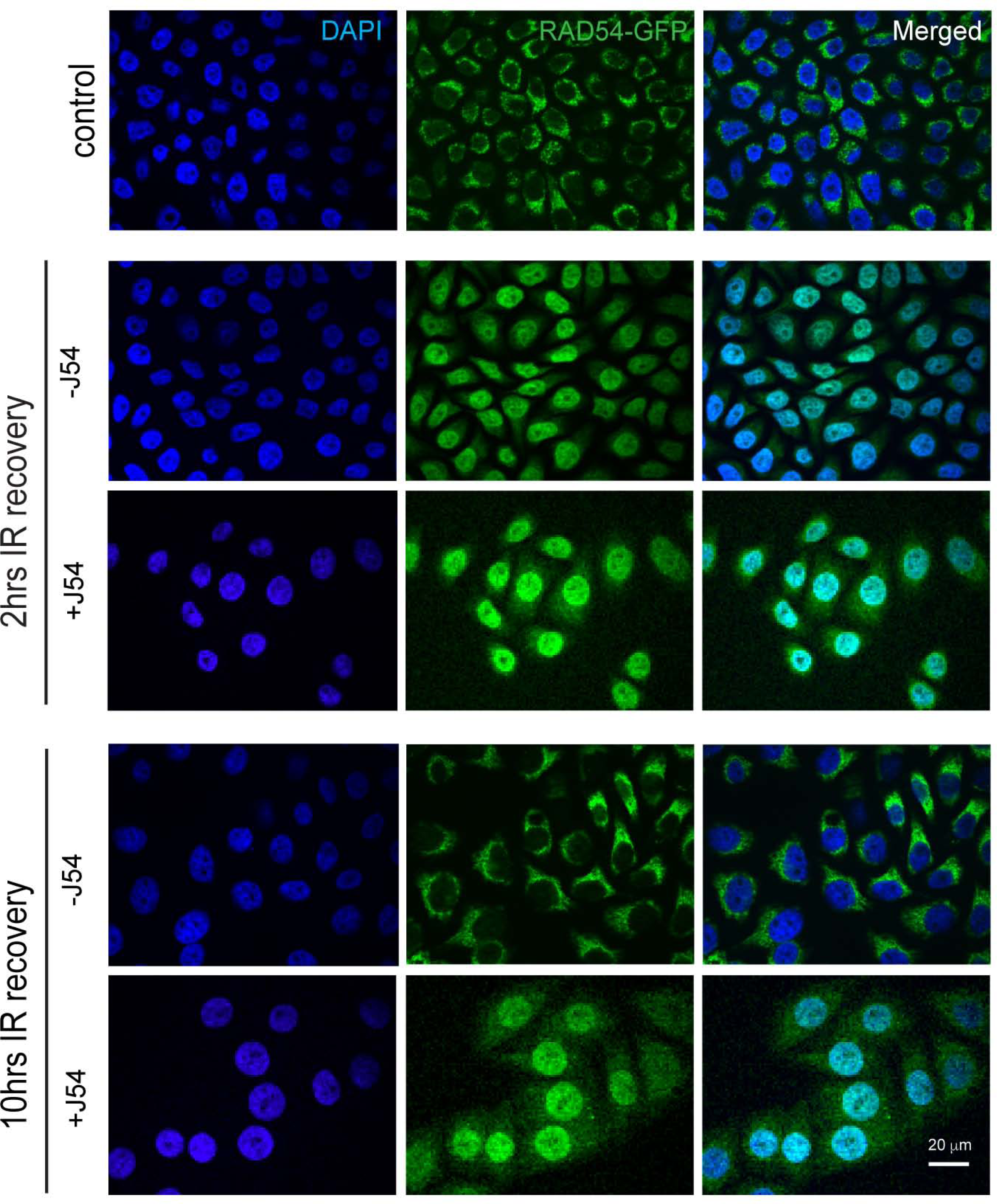
RAD54-GFP localization changes with TLK1 inhibition after IR (10Gy). At 2hrs IR recovery, RAD54-GFP is predominantly in nucleus (middle panel) and at 10hrs after IR (lower panel) when HRR attenuates, RAD54-GFP re-localizes to cytoplasm. With TLK1 inhibition followed by IR and recovery for 2hrs (2hr, +J54), RAD54GFP localizes in nucleus. At increased time of IR recovery (10hr, +J54) with TLK1 inhibition, RAD54GFP re-localization to cytoplasm is hindered. Control panel shows RAD54-GFP localization in non-irradiated cells. Images acquired from fixed cell microscopy with DAPI stained nucleus. Scale bar is 20μm.

### TLK1 mediated RAD54-phosphorylation regulates RAD51-RAD54 interaction

Phosphorylation of RAD54 at its NTD (T41, T59) or CTD (T700) may alter its affinity to RAD51(57). First, we tested if the phosphomimic RAD54 shows altered affinity to RAD51. Our results show that RAD54-T700D interacts with RAD51 more avidly than any of the other RAD54 mutants or WT protein (Figure 5A, B; repeated in Figure 5C, D), whereas the NTD phosphomimics displayed reduced affinity. As an endeavor to analyze functional interaction between the RAD54 phosphomimic mutants and RAD51, we used an ATP hydrolysis assay because it has been known that RAD51 stimulates RAD54’s ATPase activity through direct interaction(58,59). First, we tested the ATPase activity of RAD54 WT, T2D, T3D (triple phosphomimic), and T700D per se at two different KCl concentrations (22.5 and 75 mM KCl). As shown in Figure 5E-F and Figure S13, the RAD54 phosphomimic proteins are similarly active in hydrolyzing ATP as the wild type protein, suggesting that phosphorylation at the three threonines does not affect RAD54’s ATPase activity. In the presence of RAD51, as consistent with the published results, the ATPase activity is greatly stimulated by RAD51 (∼2-fold at 22.5 mM KCl, ∼15-fold at 75 mM KCl, Figure S13 and Figure 5F). The differences between RAD54 WT and RAD54 mutant proteins is clearly detectable at 75 mM KCl (Figure 5F). Compared to stimulation of ATPase activity in RAD54-WT by RAD51, phosphomimic mutants in NTD (T2D and T3D) shows reduced ATPase activity and importantly RAD54-T3D exhibits significantly reduced ATPase activity (∼50% less) than the wild type protein (Figure 5E and F). Collectively, these results show that the RAD54 mutations do not affect the intrinsic RAD54 ATPase, while the phosphomimic mutations (T2D and T3D) compromise the stimulation effect of RAD51 and thus decrease the ATPase activity of the RAD51-RAD54 complex under more physiological ionic strength conditions.

**Figure 5:**
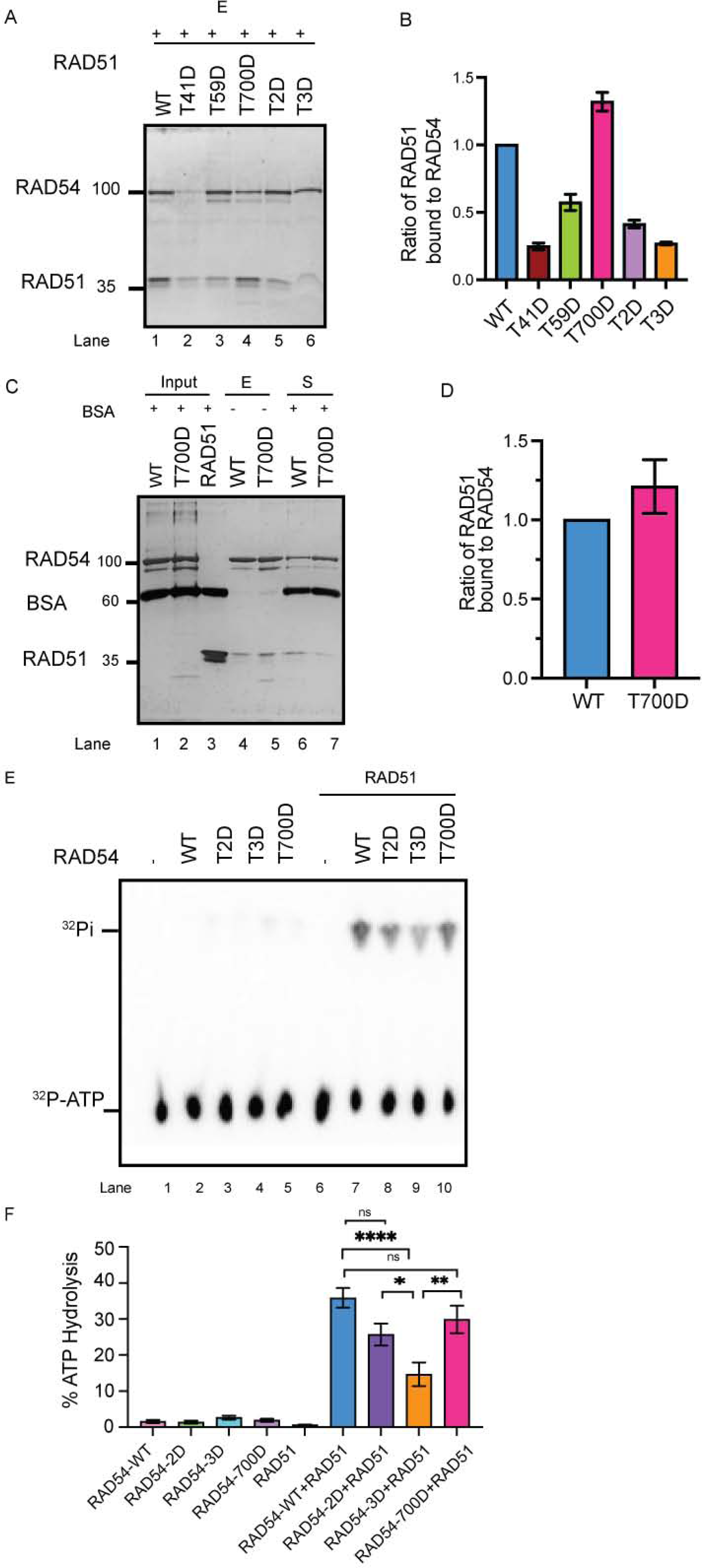
RAD54-T700D binds RAD51 with higher affinity. A-B) RAD54 phosphomimic mutant proteins exhibit different affinities to RAD51 than RAD54-WT (reactions eluted from beads shown in silver-stained gel in A and quantified in B). C-D) RAD54-T700D shows enhanced affinity for RAD51 (Eluate (E) and Supernatant (S) fraction shown in silver-stained gel (C), and quantification shown in (D). E-F) ATPase activity of RAD54-WT, NTD mutants (T2D, T3D) and CTD mutant (T700D) with dsDNA shown as %ATP hydrolysis (y-axis) in TLC and in the absence and presence of RAD51(E). Quantification of the results (F). One-Way ANOVA, Tukey’s multiple comparison test (*, p<0.05; **, p<0.01; ****, p<0.0001).

We tested whether chemical inhibition of TLK1 with J54 affects the association of RAD54 with RAD51 in cells after 24hrs treatment. It is important to note that we have a phosphor-specific antibody available for the T700 only, a residue that is phosphorylated during recovery from IR and during later stages in the HRR process. Thus, we do not know the timing of phosphorylation at the two NTD residues or whether it is induced after IR. We determined the fraction of RAD51 associated with RAD54 in unperturbed condition (Figure S14) and although RAD54 was quantitively bound to the HA beads, very little of the available RAD51 was retained with it, suggesting that without induced DNA damage RAD54 and RAD51 are not readily found in a complex. However, after treatment with J54, significantly more RAD51 co-purifies with RAD54 suggesting that the phosphorylation of RAD54 by TLK1 acts to impede the association of the two proteins under unperturbed conditions.

### Phosphomutant RAD54-T700D delays HRR kinetics

Although HRR occurs during S/G2 phase, HeLa cells constitutively express RAD54 and RAD51 and the expression increases post DSB induction(60,61). Further, RAD54 and RAD51 co-localize upon exposure of cell to IR(18,51). We wanted to test if RAD54 and RAD51 foci co-localize upon exposure of cells to DSB inducing drug, Mitomycin C (MMC). Here, we treated HeLa cells with MMC (3 μM) for 2hrs and used immunocytochemistry (ICC) to visualize RAD54 and RAD51. We find that 4hrs post MMC induction, the number of RAD51 foci/nucleus increases, indicative of ongoing HRR, and that RAD51 and RAD54 foci strongly correlate (Figure 6A-C). A similar observation was made for irradiated cells (Figure S15)

**Figure 6:**
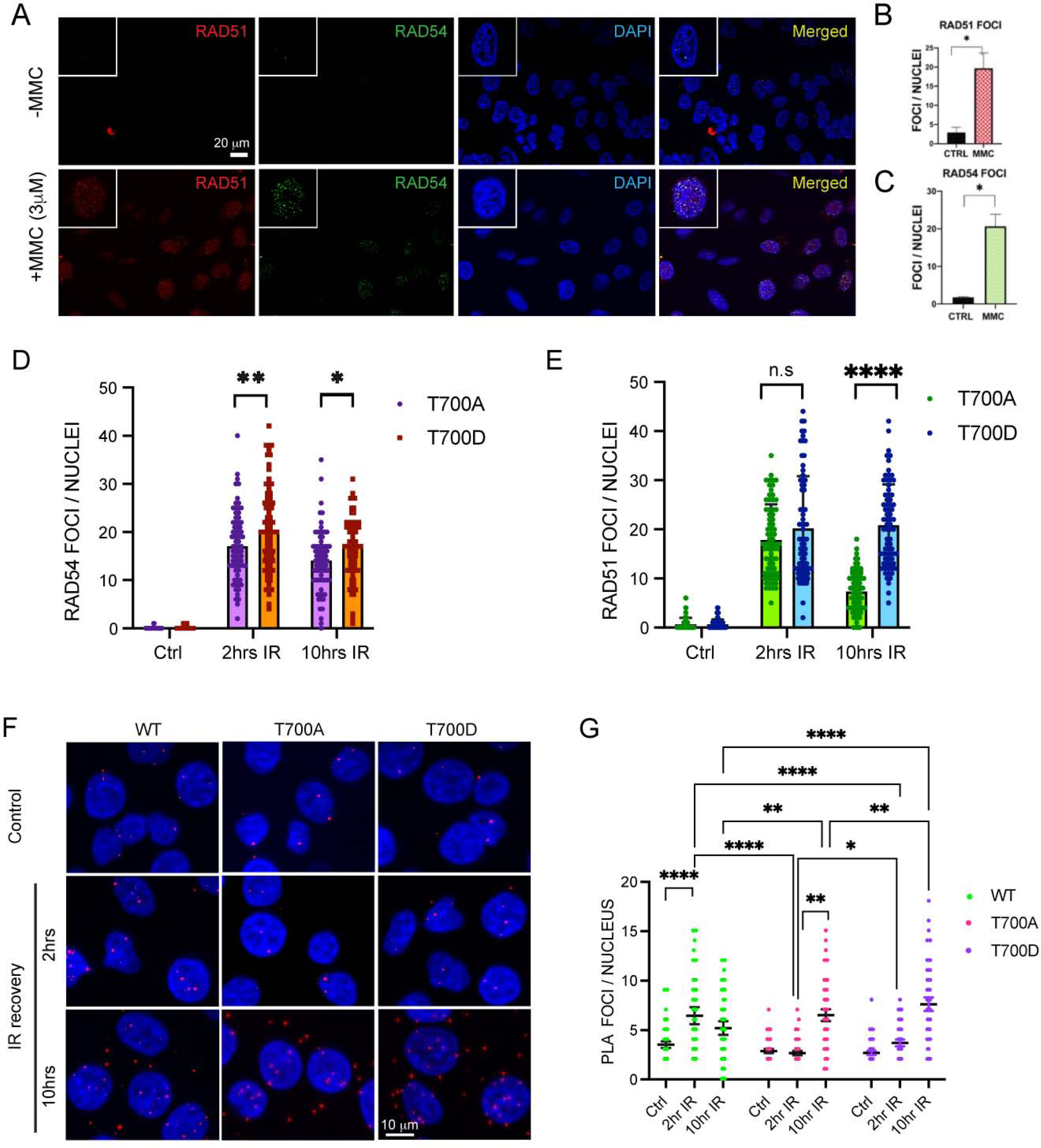
Phosphorylation at RAD54-CTD (T700) leads to delayed HRR kinetics. A-C) RAD54 and RAD51 foci increase upon exposure of cells to MMC HeLa cells were treated with MMC and allowed to recover for 4hrs, fixed and stained (RAD51: red, RAD54: green) cells shown in panel A. Quantitation of RAD54 foci/nucleus (shown in panel B) and RAD51 foci/ nucleus (shown in panel C). Total of 90-100 nuclei counted, the results are mean ± SEM from three biological repeats, with p-values (*, p<0.05) determined by Unpaired Student’s t-tests with Welch’s correction. Scale bar 20μm. D-E) RAD54 and RAD51 foci formation in HeLaRAD54KO cells expressing RAD54-T700A or RAD54-T700D. Cells were exposed to 10 Gy X-rays and allowed to recover for 2 and 10 hrs. 70-80 nuclei were assessed from 3 independent experiments. Statistical significance test performed with 2Way ANOVA, with Šídák’s multiple comparisons test), (*, p<0.05; **, p<0.01; ****, p<0.0001). F-G) Results from WT, RAD54-T700A and RAD54-T700D interaction with RAD51 in PLA assay probed with anti-RAD54 and anti-RAD51. (F) Representative micrographs; scale bar: 10μm. G) Quantitation of the nuclear PLA signals from 150 cells were assessed from 3 independent experiments. Statistical significance test performed with 2Way ANOVA, multiple comparison (simple effects within rows), (*, p<0.05; **, p<0.01; ****, p<0.0001). The specificity of the RAD54/51 PLA interaction was previously verified using the RAD54-KO cells for negative control (51).

Since purified RAD54-T700D shows enhanced affinity to RAD51 as compared to the WT protein, we wanted to test the co-distribution of RAD54-T700A/D with RAD51 in cells. We induced DNA damage with 10 Gy IR and allowed cells to recover for 2hrs and 10hrs. We find that, RAD54-T700A forms significantly fewer RAD54 foci than T700D during initial and later IR recovery (Figure 6D and Figure S16). Interestingly, RAD54-T700A and T700D showed similar RAD51 foci count at initial IR recovery (2hrs) while at later IR recovery (10hrs) T700A had significantly less RAD51 foci than T700D (Figure 6E and Figure S16). Next, we wanted to examine the RAD54-T700A/D-RAD51 interaction in cells. We performed the Proximity Ligation assay (PLA) in fixed cells using RAD54 and RAD51 antibody, where a signal or PLA focus indicates association of RAD54 and RAD51 within ∼ a 40 nm range (62,63). At 2hrs post IR, RAD54-WT cells showed the highest number of nuclear PLA foci, and RAD54-T700D cells showed significantly higher numbers of nuclear PLA foci than RAD54-T700A cells, although PLA foci are lower in RAD54-T700D than in WT cells (Figure 6F middle panel and quantification in Figure 6G). These results suggest that at 2hrs post IR, phosphorylated RAD54-T700 may interact more tightly with RAD51 than unphosphorylated RAD54-T700. We wanted to study if the interaction between RAD54 and RAD51 persists at later HRR stage (10hrs post induction). We observed that while WT cells have a decrease in number of PLA foci, T700A and T700D had increased foci. Importantly, T700D had significantly higher interaction (number of foci) than T700A (Figure 6F, bottom panel and quantification in Figure 6G). The specificity of PLA signal was confirmed by no primary antibody controls (Figure S17). We confirmed the RAD54-T700A/D interaction with RAD51 for the later time-point through immuno-precipitation experiments (Figure S18A-B). From these findings we conclude that T700D has higher association with RAD51 during recovery from IR. At a later stage (10hrs), however, complex formation between RAD54-T700D and RAD51 may be inhibitory to HRR completion, possibly delaying branch migration or DNA polymerase extension.

### Molecular modelling suggests RAD54-T700 lies in proximity to RAD51

The RAD54-T700 (bold) residue lies within the peptide SDC**T**SDLAG in a unique site for two reasons, firstly it has two negatively charged residues (Asp acid) at the −2 and +2 positions, which makes the environment ionic and hence possibly solvent accessible. Secondly, the Zn-finger like motif (H-676, C-681, C-684, H-708) which contacts the DNA backbone surrounds T-700 residue within the CTD (29). We showed that T700D has strong association with RAD51. Therefore, we speculate that T700 phosphorylation may create an ionic interaction and, as such, affect the interaction of RAD54 with dsDNA and RAD51.

The limited availability of structural data for the human RAD54 protein led us to model the structures of RAD54-WT and the RAD54-T700D mutant protein using complete sequence. We modelled both protein structures RAD54-WT and RAD54-T700D by method as described above (Figure S19 and S20). The structural assessment of modelled RAD54-WT protein showed a MolProbity score of 1.72 and Qmean Score of −2.98, visual inspection of the model shows the N-terminal region to be unstructured (Figure S19C, E). This model was subjected to MDS for 500 ns, the simulation trajectory was analysed to understand the structural changes occurring during the simulation. The RMSD of the protein initially converged slowly between 5-7 Å during initial period of simulation, later it stabilised between 3-4 Å for 200 ns. A slight steep rise in the RMSD was observed around 270 ns with a slight rise in the RMSD, later the RMSD stabilises between 1-2 Å for rest of the simulation suggesting the stability of the protein which also indicates that the protein might have achieved a conformation stability, this was ascertained by the visual inspection of the MD trajectory. The N-terminal gets into some more stable conformation and stay in the same form for about 200 ns with minimal fluctuation.,. These changes are also reflected by the RMSF plot showing high fluctuations as high as 20 Å for the residues belonging to the NTD (Figure S19B). A representative frame from end of the simulation trajectory was extracted and analysed, it showed MolProbity score of 1.55 and Qmean Score of −2.52, which is better than the initial scores, lower these scores better are the structural features and stability of the protein conformation (Figure S19D, F). The structural assessment of modelled RAD54-T700D protein showed a MolProbity score of 1.45 and Qmean Score of −2.90, visual inspection of the model shows the N-terminal region to be unstructured as in case of the WT (Figure S20C, E). This model was subjected to MDS for 500 ns, the simulation trajectory was analysed to understand the structural changes occurring during the simulation. The RMSD of the protein initially converged slowly between 1-7.5 Å during first 200 ns of the simulation. After 200 ns the trajectory shows high fluctuations for about 100 ns during which the RMSD fluctuated between 2.5-4 Å, after which it remained between 2-3 Å for rest of the simulation time. The visual inspection showed the structural changes in the N-terminal domain during the 200-300 ns which later stabilised indicating that the protein might have achieved a conformation stability (Figure S20A). These changes are also reflected by the RMSF plot showing high fluctuations as high as 12 Å for the residues belonging to the NTD (Figure S20B). A representative frame from end of the simulation trajectory was extracted and analysed, it showed MolProbity score of 1.56 and Qmean Score of −2.47, which is better than the initial scores, lower these scores better are the structural features and stability of the protein conformation (Figure S20D, F). Along with RAD54, structural data of RAD51 is also limited to some extent so we modelled the structure of RAD51 protein as well. The modelling was performed using two different tools, first we used AlfaFold (https://alphafold.ebi.ac.uk/)(36) and second one was the CCG MOE(37). The models obtained for these tools were compared, ranked and superposed to ascertain the quality of generated models. In both cases the models had some unstructured regions especially on the terminal regions. These models were refined by subjecting them to the molecular dynamics simulation (MDS) for 500ns using explicit solvent conditions. The RAD51 sequence consisted of 339 residues, these were modelled and the resultant model was analysed for its structural integrity (Figure S21A-Fa-f). The structural assessment was performed by the SWISS-MODEL structural assessment tool where, Ramachandran analysis, MolProbity scoring and Q-mean score analysis was performed(38,64). The modelled RAD51 shows an unstructured N-terminal (Figure S21Cc), its MolProbity score was 1.41 and the Q-mean score was −1.59. This model was subjected to MDS for 500ns in explicit solvent model. The MDS trajectory was analysed with CPPTRAJ(47) and representative structure towards the end of the simulation was extracted. The trajectory analysis shows initial fluctuation of around 2-3 Å at the beginning of the simulation, these fluctuations (RMSF) stabilised to around 2 Å between 100-200ns (Figure S21Bb). These RMSD fluctuations again appear between 4-5 Å for 200 to 300 ns suggesting the folding of the N-terminal into a more stable conformation, this can be observed from the RMSF plot, where it resides between Met1 to Thr90 shows higher fluctuations (Figure S21Aa). The extracted frame towards the end of the simulation was again subjected to structural assessment. It showed a reduced MolProbity score of 1.27 and the Q-mean score was −1.54 suggesting a better stabilisation of the modelled RAD51 (S21Dd).

To understand the interaction between the RAD54 and RAD51 and the effect of mutation in RAD54 on this RAD54-RAD51 interaction we performed docking studies with Haddock protein-protein docking tools. Several different docking experiments were performed for RAD54-WT-RAD51 and RAD54-T700D-RAD51 complexes with the MOE protein docking protocol, the standard docking protocol with flexible docking interface was performed with several rounds of different combinations and the models were ranked based on the dock score and Z-score (Table 1). The docking results were compared for both the tools and a consensus was generated to narrow down with the protein-protein interaction complex for the RAD54-WT-RAD51 and RAD54-T700D-RAD51 complex. The best complex for RAD54-WT-RAD51 showed a dock score of −88.7 (5.3) and Z-score of −1.6, whereas for the RAD54-T700D-RAD51 showed a dock score of −99.9 (6.4) and Z-score of −1.4. The results obtained from the Haddock docking were further investigated by MDS for 500 ns explicit solvent model. The RAD54-WT-RAD51 showed major fluctuation in the simulation of about 4 Å between 125-250 ns, but this became more stable towards the rest half of the simulation, it showed the RMSD of about 2.0 Å in the last 240 ns of the simulation (Figure 7A). The RMSF of this complex shows the high RMSF for the residues belonging to the N-terminal of the RAD54-WT in the range of 7.5 Å for the residues in the terminal region, the sequence of RAD54-WT consisted of 747 residues. Similarly, we observed higher fluctuation in the RMSF of the RAD51 in the N-terminal and C-terminal domains of about 5 Å (Figure 7D). The structural and trajectory visualisation shows that RAD51 N-terminal segment interact with the RAD54 C-terminal section during the initial phase of the simulation which further shifts slightly towards the residues in the range of Arg683-Ser701 (Figure 7E, F). In order to further investigate these interactions, we performed a distance-based hydrogen bond analysis for the selected residues at the interaction interface of RAD54-WT-RAD51 complex. The distance-based hydrogen bond analysis shows the interaction between several residues during the period of 130-140 ns these were in the range of 3.5 Å distance from each other, this is also supported by the RMSD plot during this period. These residues again interacted to form the hydrogen bond interactions towards the end of the simulation as shown in the plot (Figure 7B, G). The RAD54-WT(Glu371)-RAD51(His244) (Black: 2.87), RAD54-WT(Arg374)-RAD51(Asp198) (Red: 2.80), RAD54-WT(Glu378)-RAD51(Lys64) (Green: 2.69 Å), RAD54-WT(Arg691)-RAD51(Glu29) (Blue: 2.80 Å). These distances were calculated between the hydrogen donor-acceptor pairs in these residues, it shows the formation of hydrogen bonds at certain instances during the MDS. These protein-protein interactions suggest for an intermittent interaction between various residues in the RAD54-WT-RAD51 complex. The principal component analysis for this complex was performed to gain insight into its various components. The PCA suggest for two different components of the Complex with a major conformational group which is away from or different from the initial state. The Black marker in the PCA plot (Figure 7C) refers to the initial structure of the complex during MDS and the Red dots shows the variance in the conformational space of the complex. The MDS trajectory for 500 ns was further calculated for extended 50 ns which was used for calculating the MMBSA of the complex. The RAD54-WT-RAD51 complex showed a ΔG_bind_ = −85.55 (7.61) Kcal/mol, various components of MM-GBSA and their contributions are provided in the Table 1.

**Figure 7:**
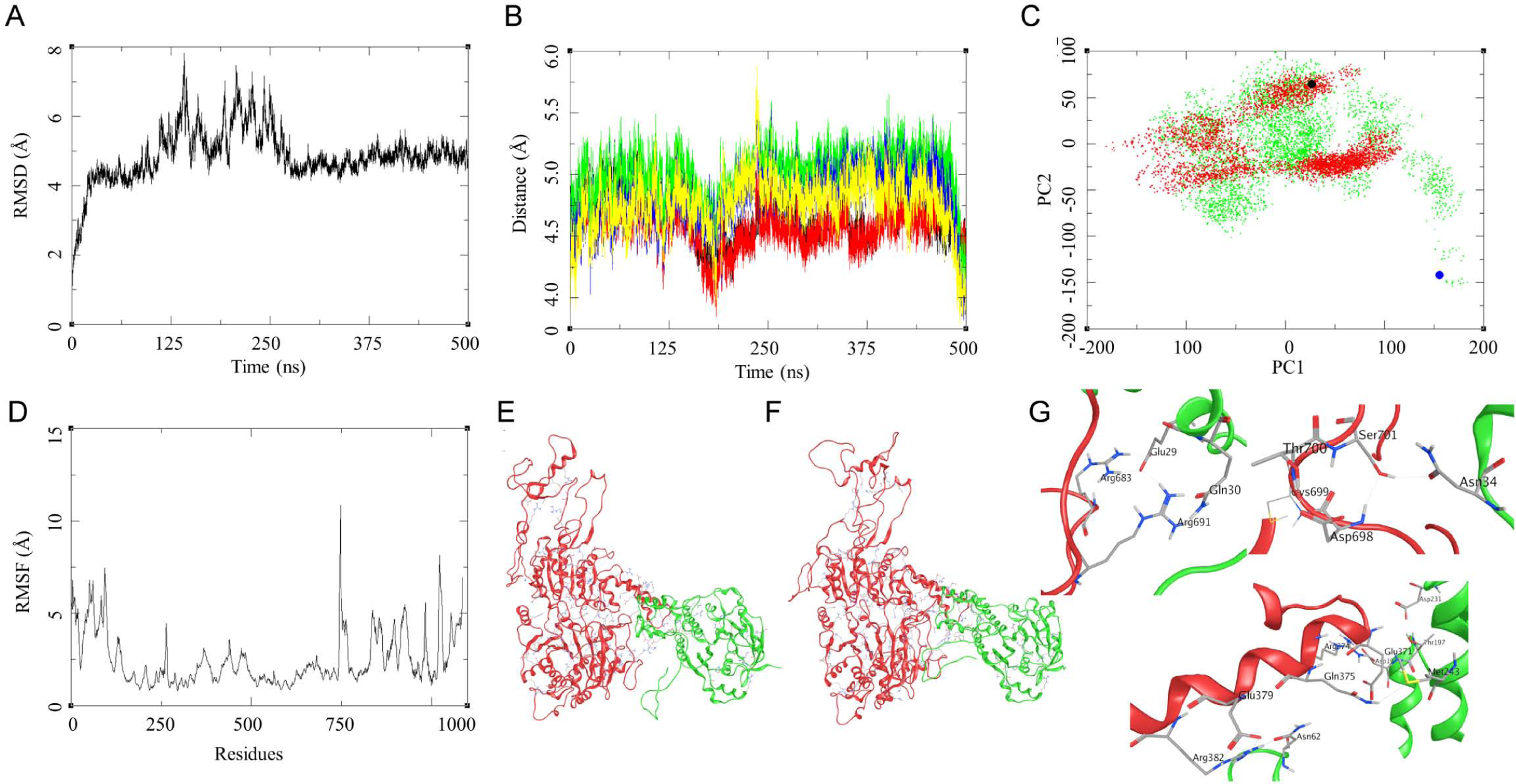
Modelling studies on the RAD51-RAD54-WT interaction. (a) RMSD of the RAD51-RAD54-WT complex over 500 ns; (b) Distance based hydrogen bond analysis: (1) GLU371-HIS244 (Black), ARG374-ASP198 (Red), GLU378-LYS64 (Green), ARG691-GLU29 (Blue); (c) Principal Component Analysis (PCA) for RAD51-RAD54-WT (Red), Black marker as the reference complex structure and RAD51-RAD54-T700D -Mutant (Green), Blue marker as the reference complex structure; (d) RMSF of the RAD51-RAD54-WT complex over 500 ns; (e) Initial state of RAD51-RAD54-WT complex at the beginning of 500 ns; (f) Final state of RAD51-RAD54-WT complex at the end of 500 ns; (g) Residue level interaction between RAD51 and the RAD54, Arg683-Glu29, Arg691-Gln30, Ser701-Asn34, Asp698-Ser701, Glu379-Asn62, Arg382-Asn62, Gln375-Met243 and Arg374-Glu37.

**Table 1.** Protein-protein docking studies and the binding free energy components for the protein-protein complexes calculated by MM-GBSA analysis, all energies are in Kcal/mol with standard deviation in parenthesis.

| Models | Haddock Docking studies |  | MD trajectory-based MM-GBSA* |  |  |  |  |  |  |
| --- | --- | --- | --- | --- | --- | --- | --- | --- | --- |
| | Dock Score | Z-Score | $\Delta E_{VDW}$ | $\Delta E_{ELE}$ | $\Delta G_{GB}$ | $\Delta G_{Surf}$ | $\Delta G_{gas}$ | $\Delta G_{Sol}$ | $\Delta G_{bind}$ |
| **RAD54-WT-RAD51** | -88.7 +/- 5.3 | -1.6 | -119.37 (7.94) | -1725.19 (46.61) | 1777.11 (46.01) | -18.10 (1.04) | -1844.56 (47.57) | 1759.01 (45.62) | -85.55 (7.61) |
| **RAD54-T700D-RAD51** | -99.9 +/- 6.4 | -1.4 | -100.45 (6.96) | -1667.35 (47.58) | 1698.34 (42.23) | -16.78 (0.65) | -1767.80 (47.08) | 1681.56 (42.01) | -86.24 (9.26) |

\* $\Delta E_{VDW}$ = van der Waals contribution from MM; $\Delta E_{ELE}$ = electrostatic energy as calculated by the MM force field; $\Delta G_{GB}$ = the electrostatic contribution to the solvation free energy calculated by GB; $\Delta G_{Surf}$ = solvent-accessible surface area; $\Delta G_{Sol}$ = solvation free energy; $\Delta G_{gas}$ = gas phase interaction energy; $\Delta G_{bind}$ = Binding free energy

Similar analysis was performed for the RAD54-T700D-RAD51 complex. The MDS trajectory RMSD shows a large fluctuation though the trajectory with a steep rise of about 5 Å during first 25 ns, later the RMSD fluctuated between 5-6 Å till 375 ns after which the RMSD started stabilising between 2-3 Å for last 50 ns (Figure 8A). The RMSF of this complex shows high fluctuations for the residues belonging to the N-terminal of the RAD54-T700D in the range of 12.5 Å for the residues in the terminal region, the sequence of RAD54-T700D consisted of 747 residues. Similarly, we observed higher fluctuation in the RMSF of the RAD51 in the N-terminal and C-terminal domains of about 10 Å (Figure 7D). The RMSD and RMSF plot suggest for a large fluctuation between the interacting proteins. The visual inspection of the trajectories shows a slight shift between the interacting surface of the proteins (Figure 8C, D). Due to this we performed the distance-based hydrogen bond analysis for two different set of residues selected from the initial and final stage of the simulation. The first set consist of residues pairs similar to the ones for the WT protein and the second set is more exclusive to the T700D mutant. The first set was analysed for their distances through the trajectory and the last frame analysis was done to find their distance towards the end of the simulation so as to establish the probability of formation of hydrogen bond interactions within the complex. The RAD54-T700D (Glu371)-RAD51(His244) (Black: 5.66), RAD54-T700D (Arg374)-RAD51(Asp198) (Red: 4.22), RAD54-T700D (Glu378)-RAD51(Lys64) (Green: 6.32 Å), RAD54-T700D (Arg691)-RAD51(Glu29) (Blue: 5.85 Å). These results suggest a shift of the interacting surface with reference to that of the WT. The another set of distance-based analysis was performed for the mutant residue T700D, the distance-based hydrogen bond analysis was performed for the RAD54-D700-RAD51(Asn34) (Black: 3.55 Å), RAD54-D700-RAD51(Asn36) (Red: 6.02 Å), RAD54-D700-RAD51(Asp37) (Green: 5.63 Å), RAD54-D700-RAD51(Lys39) (Blue: 5.41), RAD54-D700-RAD51(Cys40) (Yellow: 5.39), and RAD54-D700-RAD51(Lys70) (Brown: 5.9 Å; side chain interaction D700-OD1—NZ-Lys70 = 2.80 Å, average calculated from the atom to atom distance through the trajectory) (Figure 8B). The mutant residue forms hydrogen bond interaction during the MDS with the Lys70 and other residues in the proximity. The PCA analysis of the trajectory shows a transition of the complex from its docked state to a more stable form, the blue marker in Figure 7C is the docked complex of RAD54-T700D-RAD51. The group of green dots explore the conformational space around the WT complex suggesting the similarity in the nature of interactions. This led to calculation of binding energy of RAD54-T700D-RAD51 complex, the ΔG_bind_ = −86.24 (9.26) Kcal/mol, which is slightly higher than the WT. This could be attributed to some of the interactions that are formed by the RAD54-T700D-RAD51 in addition to the RAD54-WT.

**Figure 8:**
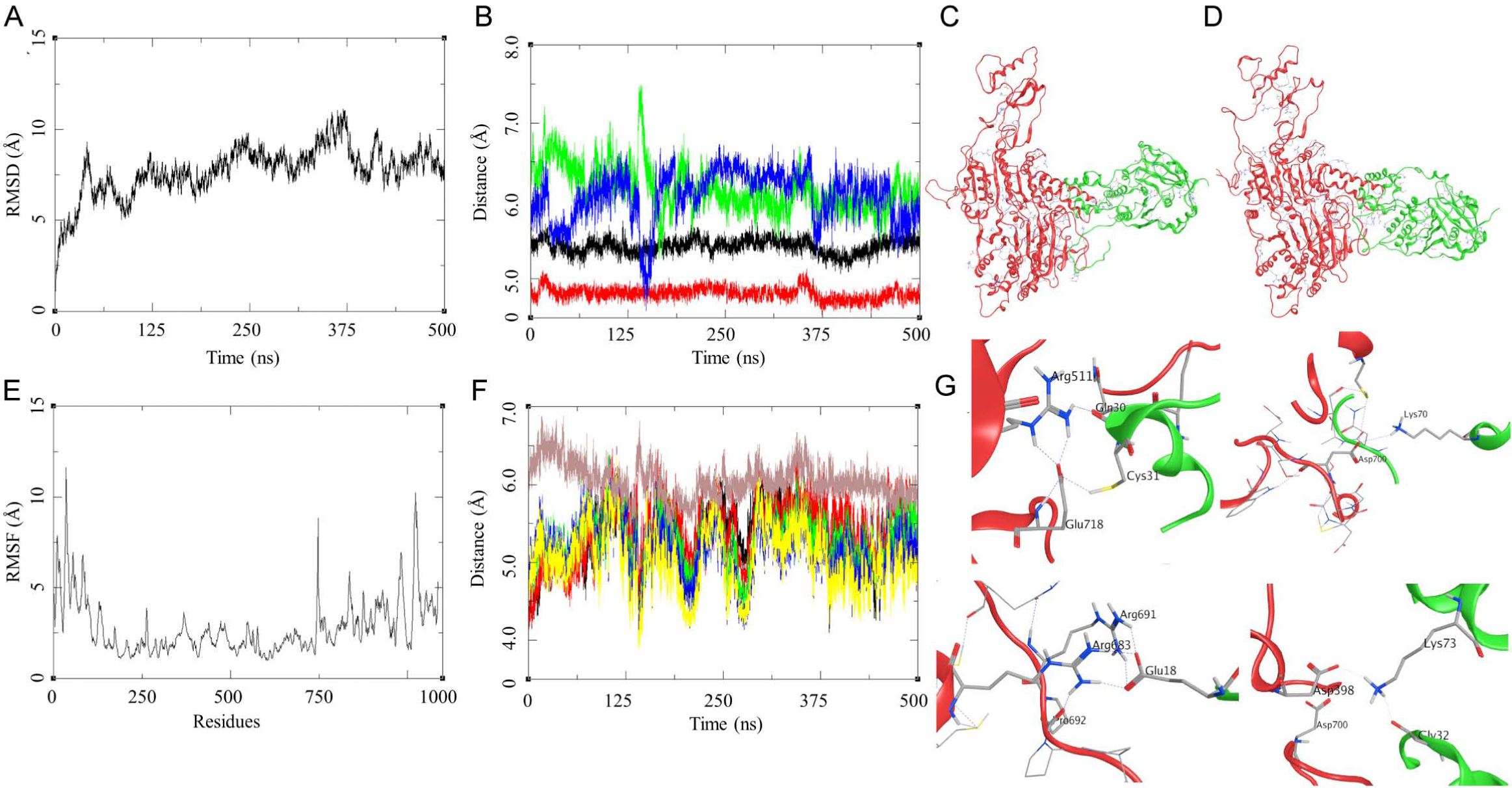
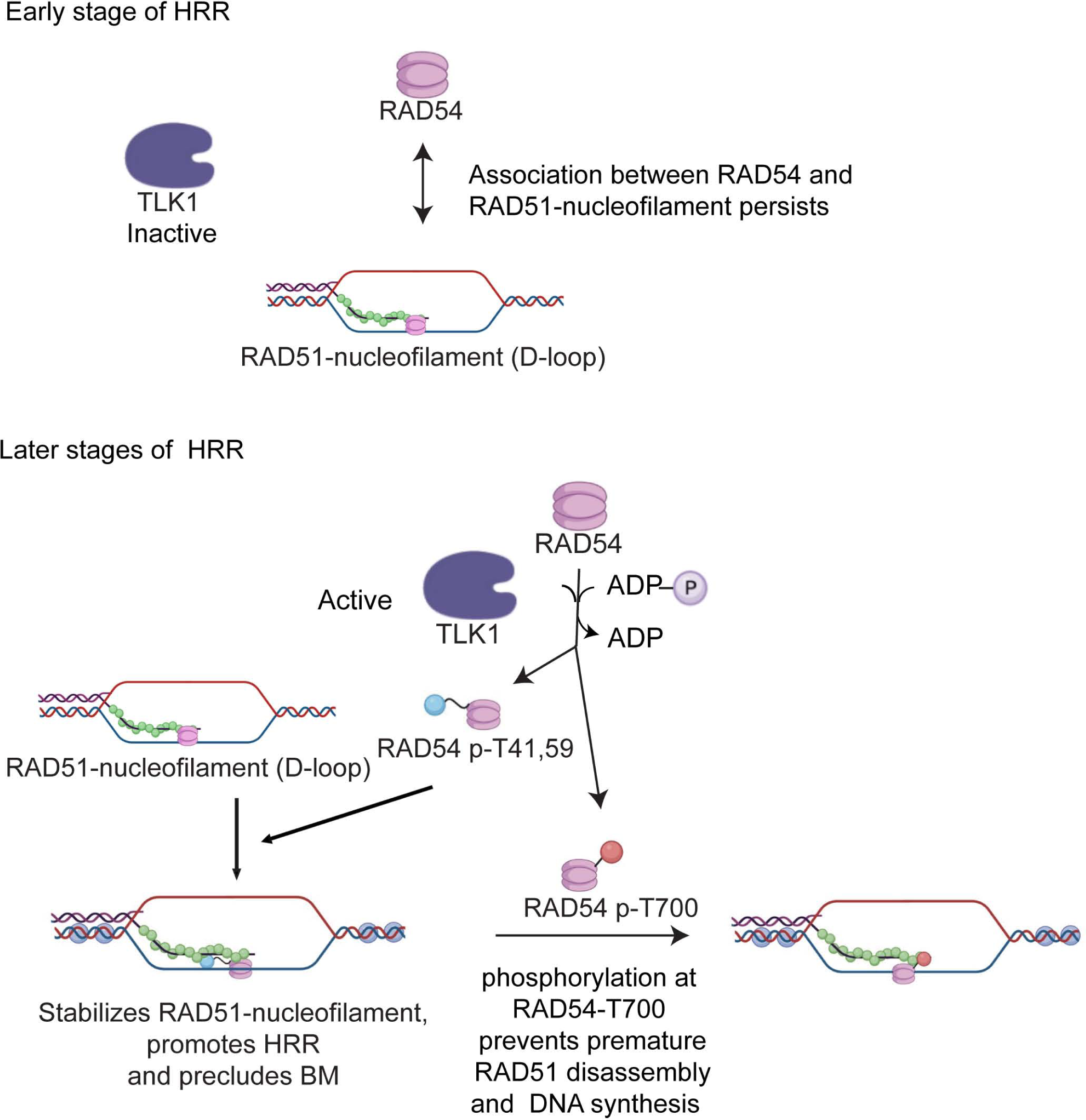
Modelling studies on the RAD51-RAD54-T700D -Mutant interaction. (a) RMSD of the RAD51-RAD54-T700D -Mutant complex over 500 ns; (b) Distance based hydrogen bond analysis: (1) GLU371-HIS244 (Black), ARG374-ASP198 (Red), GLU378-LYS64 (Green), ARG691-GLU29 (Blue); (c) Initial state of RAD51-RAD54-T700D complex at the beginning of 500 ns; (d) Final state of RAD51-RAD54-T700D complex at the end of 500 ns; (e) RMSF of the RAD51-RAD54-T700D complex over 500 ns; (f) Distance based hydrogen bond analysis: (1) D700-ASN34 (Black), D700-ASN36 (Red), D700-ASP37 (Green), D700-LYS39 (Blue), D700-CYS40 (Yellow), and D700-Lys70 (Brown) (g) Residue level interaction between RAD51 and the RAD54-T700D, Arg511-Gln30, Glu718-Cys31, Asp700-Lys70, Arg691-Glu18, Arg683-Glu18, and Asp398-Lys73.

## Discussion

Studies have shown that lack of TLK activity leads to replication fork stalling and the accumulation of single-stranded DNA. Thus, exhibiting synthetic lethality with PARP inhibitors (5,65). TLK1 has been shown to regulate the chromatin compaction state (66). One should keep in mind that, earlier studies with TLK1 shows that TLKs are involved in the DDR and/or repair following the observation that activity of TLKs is initially inhibited after IR(67,68). This inactivity could result in the rapid dephosphorylation of the RAD54-NTD residues, enabling association with RAD51 and co-translocation to the nuclei. Interestingly, phosphorylation of T41 also was identified in a phospho-proteome screening of RIP3-dependent phosphorylation events in mouse embryonic fibroblasts(69), although the responsible kinase and the significance of T41 phosphorylation has remained unknown. Although we don’t have antibodies to pT41 and pT59 that can confirm this, it is notable that in Figure 3 the pT700 phosphorylation (TLK1 dependent), while already low in untreated cells, initially disappears at 0.5h after IR, and only increases at later recovery times. It is tempting to speculate that there is a sequential pattern of TLK1-dependent RAD54 phosphorylation/de-phosphorylation during the progression of HRR.

A main observation from our studies is that TLK1 phosphorylates RAD54-T700 and, thereby, negatively regulates completion of HRR. In contrast, dual negative charge at its N-terminus positively regulates HRR, possibly by enabling RAD54/RAD51 interaction. There are several possible explanations to interpret the reason for negative regulation by T700 phosphorylation. Our preferred view is that persistent DNA damage may lead to prolonged T700 phosphorylation (as in Control of Figure 3) to avoid hyper-recombination. Another speculation is if the lesion is difficult to be repaired by HRR then phosphorylation at T700 may form the decision-making point to switch to an alternative form of non-homologous end joining (NHEJ), possibly involving the nucleolytic activity of ARTEMIS(70). It is also possible that T700D (phosphorylated RAD54-T700) prolongs the interaction between nuclear Lamin B1 and RAD51 which further stabilizes RAD51 foci post-irradiation(61). A de-phosphorylation event at T700 may occur to turn down control of branch migration and strand polymerization which may be required to allow completion of HRR. RAD54 and RAD51 appear to form spontaneous PLA foci in cells, as observed also in cytoplasmic foci with average one focus per cell. Interestingly, in vitro association of RAD51 with recombinant RAD54-T2D was reduced compared to WT protein. In contrast, we find that association of RAD51 with the pT700 is enhanced and propose that may only happen during later stage of HRR, which may be at branch migration or to prevent premature RAD51 disassembly from the filament preceding the DNA polymerization step. We speculate that RAD54 phosphorylation at T700 may also regulate the post-synapsis stage of HRR. PCNA and DNA Pol recruitment at plectonemic structures during post-synapsis stage leads to DNA dependent synthesis. Studies have shown that RAD54 depletion does not prevent PCNA recruitment(71). Rather presence of RAD54 at D-loop may halt the PCNA recruitment and phosphorylated RAD54 can further delay this process (71). In addition to maintaining genomic stability by HRR, some factors can lead to deleterious effects by HRR such as hyper-recombination leading to extensive loss of heterozygosity. Therefore, regulation of HRR serves as critical nexus to prevent tumorigenesis(2,72). We speculate that phosphorylated RAD54-T700 may complex with some unknown factors that prevent hyper-recombination or to extend the timing of RAD51 filament dissolution before HRR completion. This is undoubtedly dictated by our newly discovered modeled interaction of RAD51-Lys70 with the RAD54-pT700 (CTD) that results in stronger association of the two proteins.

Our study further opens a therapeutic axis in several HRR proficient cancer models where in Ovarian cancer (50 patients among 584 cases, https://www.cbioportal.org/study/summary?id=ov_tcga), breast cancer (20 patients among 2173 cases, https://www.cbioportal.org/study/summary?id=brca_metabric) and castration resistant prostate cancer(73)RAD54 expression is increased due to gene amplification. Targeting the activation of TLK1 dependent RAD54 phosphorylation at T700 may provide better survival strategy by downregulating HRR, based on precision medicine profile of an individual.

The lack of structural data for RAD54 and RAD54T700D and the interaction between RAD51 and its mutant led us to use the Artificial Intelligence (AI) based tool AlfaFold which assisted in building the model for the proteins. These models were validated, equilibrated and optimized for their structural features and further the Protein-protein interactions (PPI) was studied. In the RAD51-RAD54-WT PPI, the RAD51 N-terminal segment interact with the RAD54 C-terminal section during the initial phase of the simulation during which the residues in the interacting surface forms hydrogen bond interaction between the residues on the interface. These interactions are made and broken through the simulation period which suggest for good interaction between the RAD51 and the RAD54-WT leading to a stable complex. In the PPI between RAD51-RAD54-T700D RAD51, the RAD51 N-terminal interacts with the C-terminal domains of RAD54-WT with the formation of several hydrogen bonded interactions. These interactions were more dynamic than the ones in case of RAD51-RAD54-WT. The interacting surface forms various interaction between the surface residues, but one of the hydrogen bond interactions was formed by the Lys70 of RAD51 and the T700D resides of the RAD54 protein. The binding energies for both the system was calculated, it suggests that the RAD51-RAD54-T770D PPI is slightly stronger than the RAD51-RAD54-WT PPI. The structural features, protein-protein interactions and the binding free energy calculations support the findings of these interactions between RAD51 and the RAD54 proteins. While this small change in free energy might not seem much, and only results in modest increase in the pulldown assay for the T700D compared to WT (Figure 5), we recognize that were such interactions static and permanent, the next process of HRR would never happen and, for example, likely not result in the observed change in ATPase as for the T3D mutant (Figure 5F). We conclude that the results from the dynamic simulations helped a great deal in explaining the results obtained in vitro and in cells.

## Data and Code availability

All the raw data and reagents produced for this paper will be freely made available to all qualified investigators. MS data were deposited in PRIDE and are available via ProteomeXchange with identifier PXD037131.

## Acknowledgements

## Supporting information

SI-TLK-RAD54

## Acknowledgements

We would like to thank the LSUHS-Ochsner Radiation Facility for helping with Radiation treatment. We would also like to thank David Custis (Flow-Cytometry Division), and Chaowei Shang (Microscope Division), Research Core Facility, LSUHS. We are thankful to Ike-Muslow Pre-doctoral Fellowship-LSUHS (Research Distinction Award), awarded to IG. RC acknowledges the use of the Computational Shared Facility at the University of Manchester. The graphical abstract and schematic were created with BioRender.com.

## Author contributions

I.G performed most of the experiments including cell biology and molecular biology experiments and analyses described in this work. Y.K and A.B.S performed and analyzed the ATPase assays. R.C carried out the molecular modeling (HMM) and MDS experiments. J.C generated the mass spectrometry work. C.W donated the HeLa-RAD54-KO cells and collaborated on the work and production of figures. A.D.B and P.S conceived and supervised the work. I.G, A.D.B and C.W wrote the manuscript, with the contribution from all authors.

