## Supplementary material for "TLK1-mediated RAD54 phosphorylation spatio-temporally regulates Homologous Recombination Repair": SI-TLK-RAD54

**Supplementary Figures**

Supplementary Figure S1

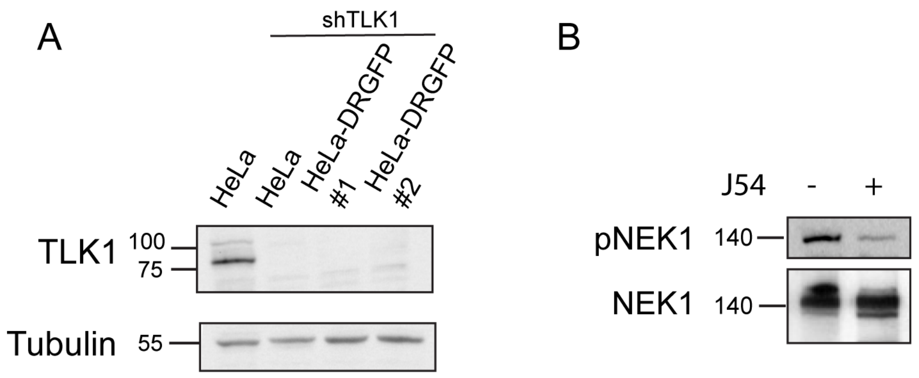


**Figure S1**: TLK1 depletion in HeLa-DRGFP cells and TLK1 inhibition in U2OS-DRGFP cells. A) Stable knockdown of TLK1 in HeLa and HeLa-DRGFP cells. shRNA against human TLK1 was transfected in HeLa and HeLa-DRGFP cells. Western blot showing TLK1 expression in HeLa cells and its absence in HeLa and HeLa-DRGFP cells (clone#1 and #2). Tubulin was used as a loading control. B) U2OS-DRGFP cells was treated with inhibitor (iTLK1, J54, 10uM). Western blot of pNek1 (TLK1 substrate) and its downregulation with TLK1 inhibitor treatment. Total NEK1 was probed using anti-NEK1.

Supplementary Figure S2

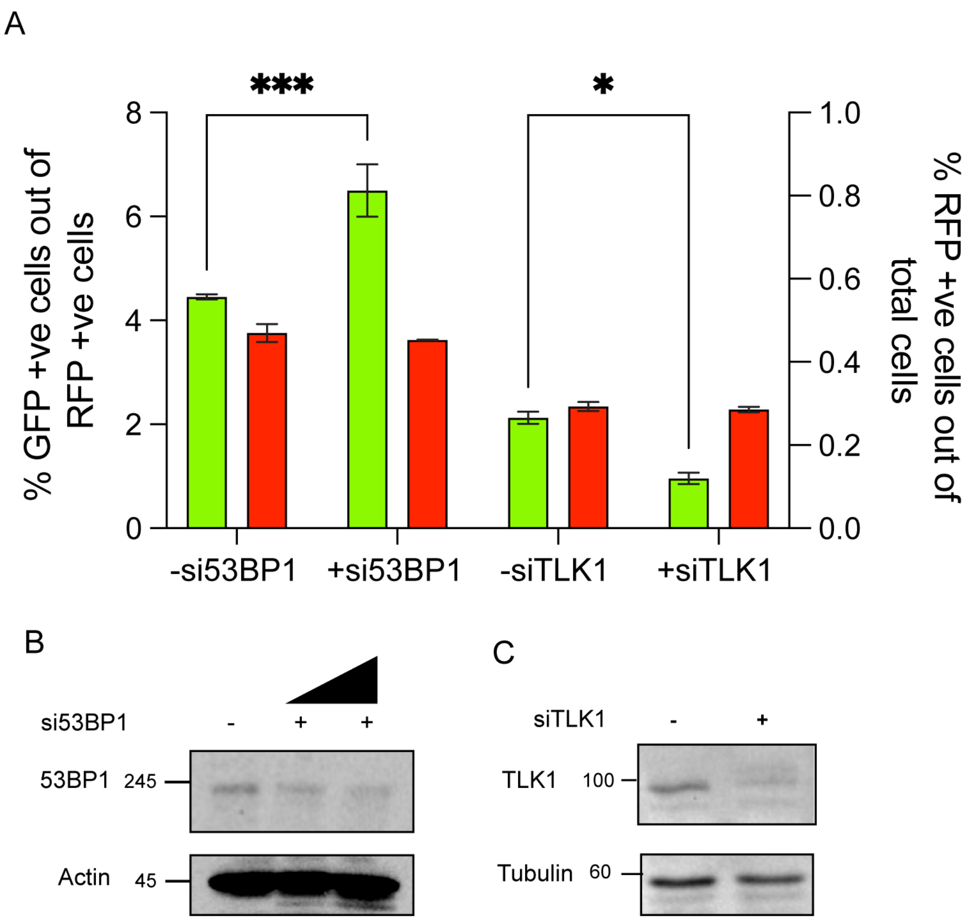


**Figure S2**: DRGFP assay in U2OS-DRGFP with depleted 53BP1 and TLK1 using siRNA. A) %GFP+ve cells and %RFP+ve cells plotted in left and right y-axis respectively in cells treated with si53BP1 (100nM) and siTLK1 (100nM). *, $p < 0.05$; ***, $p < 0.001$; 2-Way ANOVA with Tukey's multiple comparison was performed (simple effect within columns). B) Western blot probed with anti-53BP1 showing 53BP1 knock down levels using different concentrations of siRNA (50 and 100nM) were used. C) Western blot probed with anti-TLK1 antibody showing TLK1 depletion using 100nM siTLK1.

Supplementary Figure S3

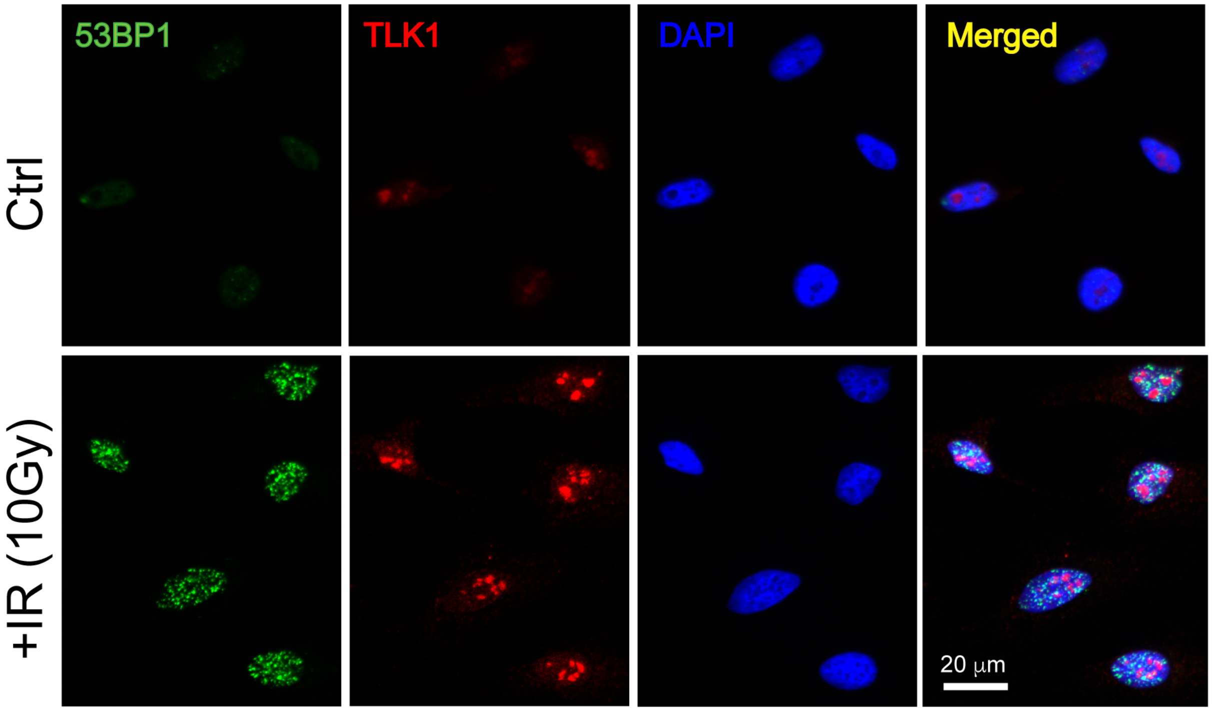


**Figure S3**: TLK1 and 53BP1 forms partly colocalized foci post-irradiation. HeLa cells were treated with 10Gy IR dose and allowed to recover for 2hrs. 53BP1 (green) and TLK1 (red) antibody used to co-stain fixed cells. DAPI stained nuclei. Scale bar is 20μm.

Supplementary Figure S4

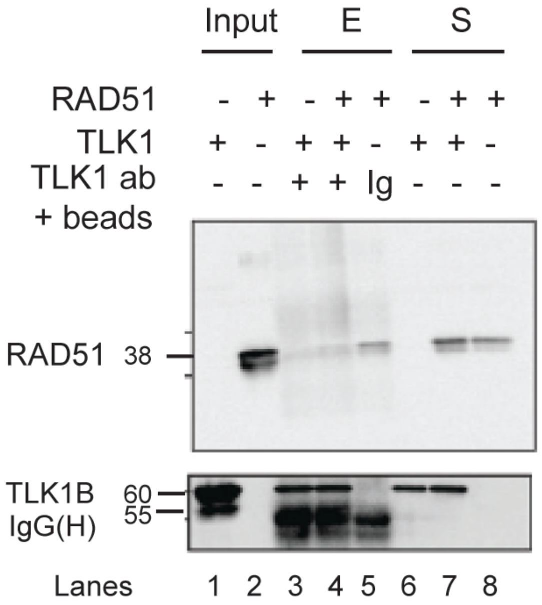


**Figure S4**: Recombinant TLK1 does not interact with RAD51 *in vitro.* Recombinant TLK1B incubated with RAD51 was immunoprecipitated using TLK1 antibody coated agarose beads. Negative control of isotype IgG (Ig) used to detect non-specific binding of proteins to antibody. Proteins eluted (E) and loaded in 10% SDS-PAGE gel for analysis by western blotting. Supernatants (S) loaded to detect unbound fraction.

Supplementary Figure S5

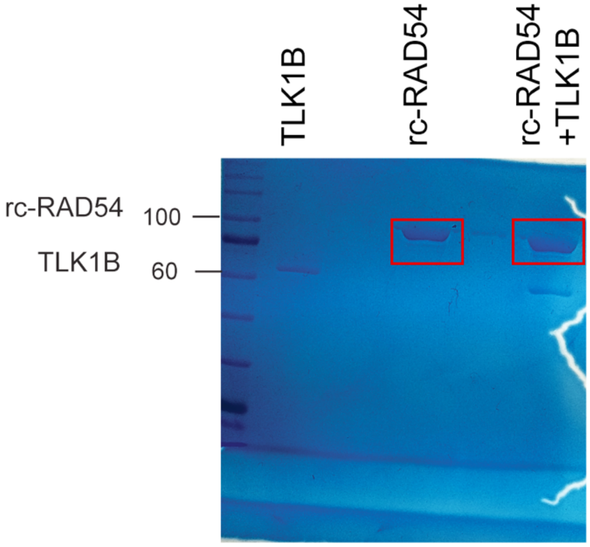


**Figure S5**: Phosphorylation of recombinant human RAD54 (rc-RAD54) by recombinant human TLK1B. Coomassie stained gel. Bands (red box) were excised and phosphopeptides analyzed by LC-MS/MS.

Supplementary Figure S6

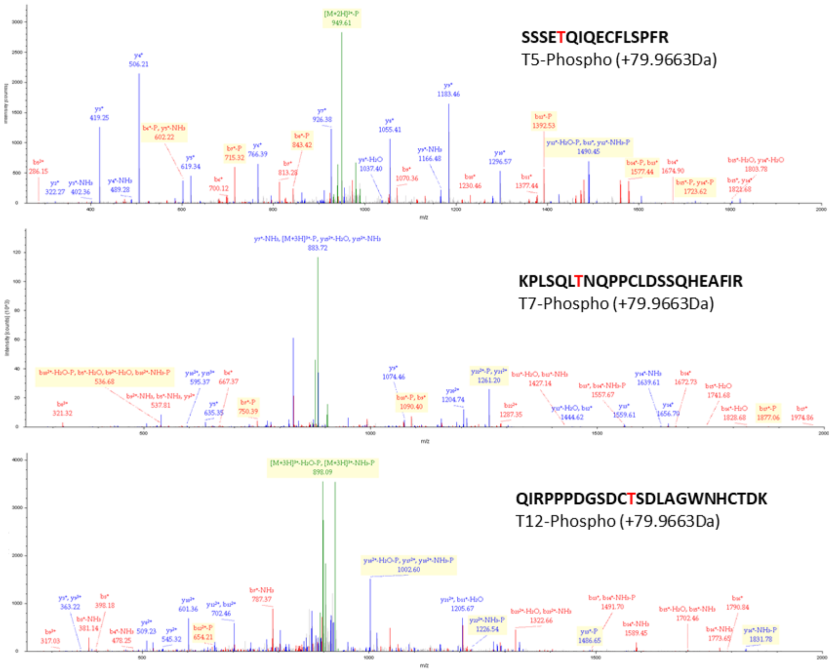


Highlighted in spectrum are detected b (red) or y (blue) ions. Green is precursor ions.

**Figure S6**: Spectral analysis of RAD54 phosphoresidue determination by MS

Supplementary Figure S7

RAD54-Control

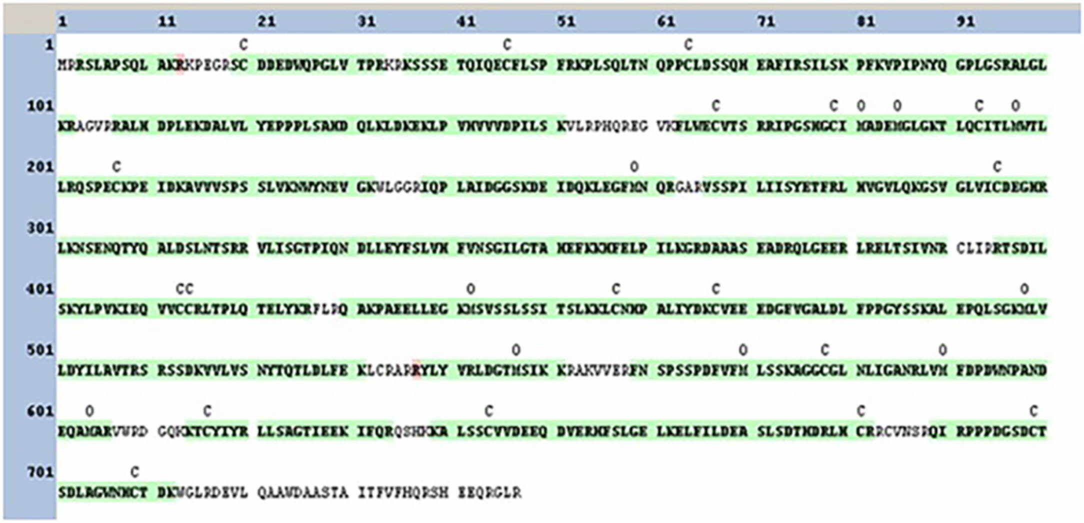


RAD54+ TLK1B

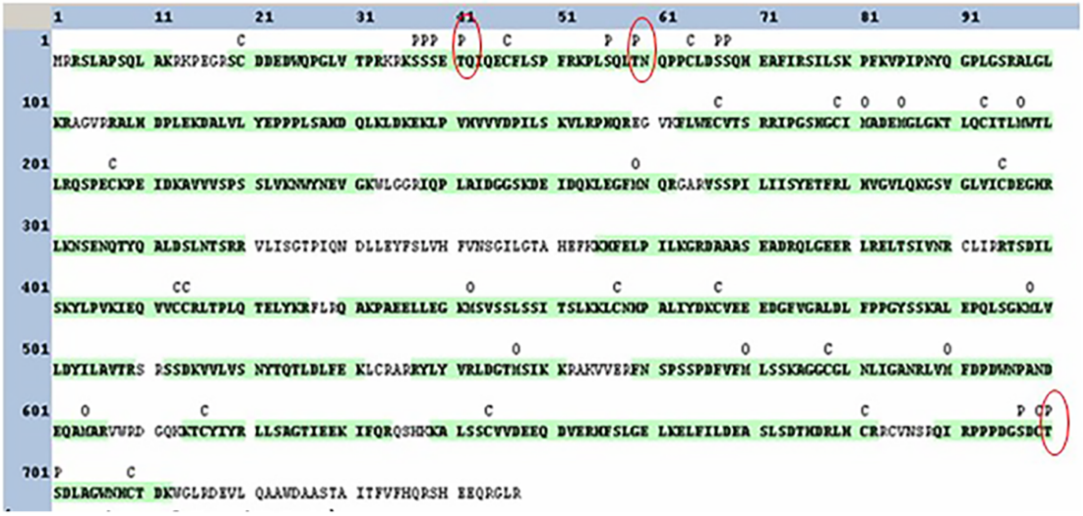


**Figure S7**: RAD54 phosphopeptide determination by MS

Supplementary Figure S8

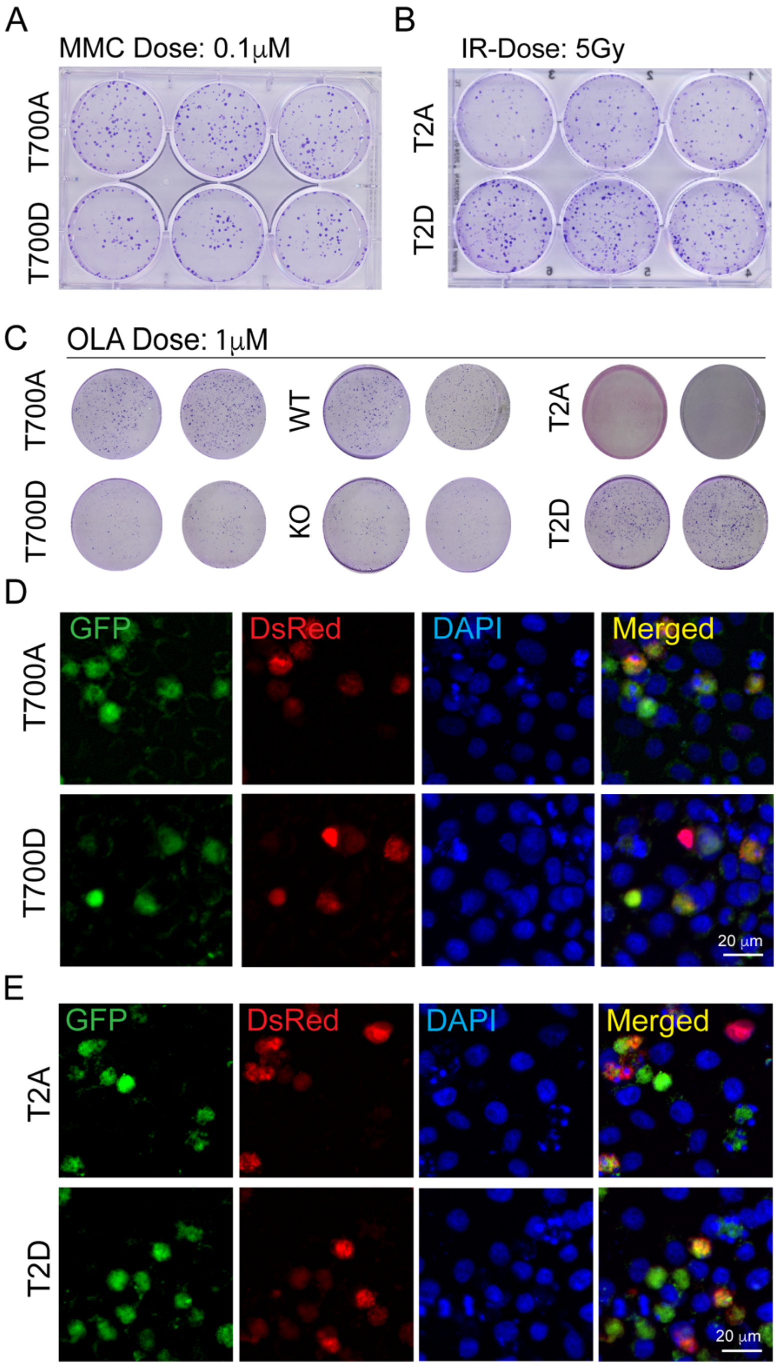


**Figure S8**: DSB sensitivity by clonogenic assay A) Representative plate of T700A/D for MMC (0.1μM), B) Representative plate of T2A/D for IR (5Gy) and C) Representative plate of T700A/D, WT, RAD54KO, T2A/D for OLA dose (1μM). D-E) HRR activity (GFP+ve and RFP+ve cells from SceI-GR-DsRed transfection) in T700A/D and T2A/D mutants. Objective 20X.

Supplementary Figure S9

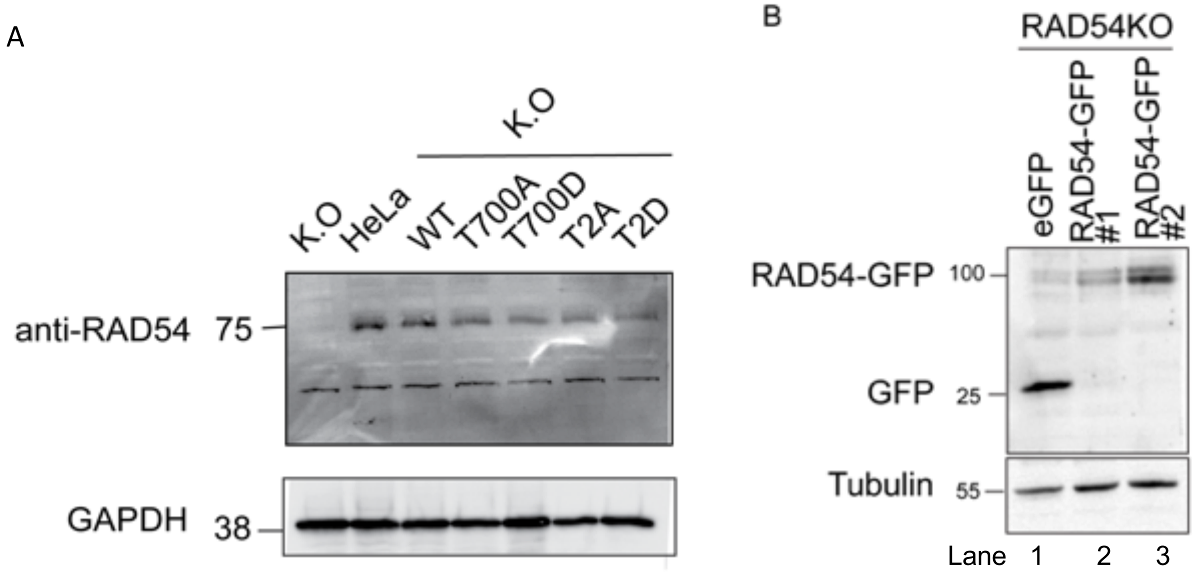


**Figure S9**: A) RAD54-WT and T700A/D and T2A/D mutant expression in RAD54KO-HeLa cells. Western blot probed with anti –RAD54 antibody. B) RAD54-GFP expression analysis in HeLa-RAD54KO cells: eGFP expressing HeLa RAD54KO used as control (lane 1) and two clones of RAD54GFP (lane 2, #1; lane 3, #2) shown in western blot probed with anti-GFP.

Supplementary Figure S10

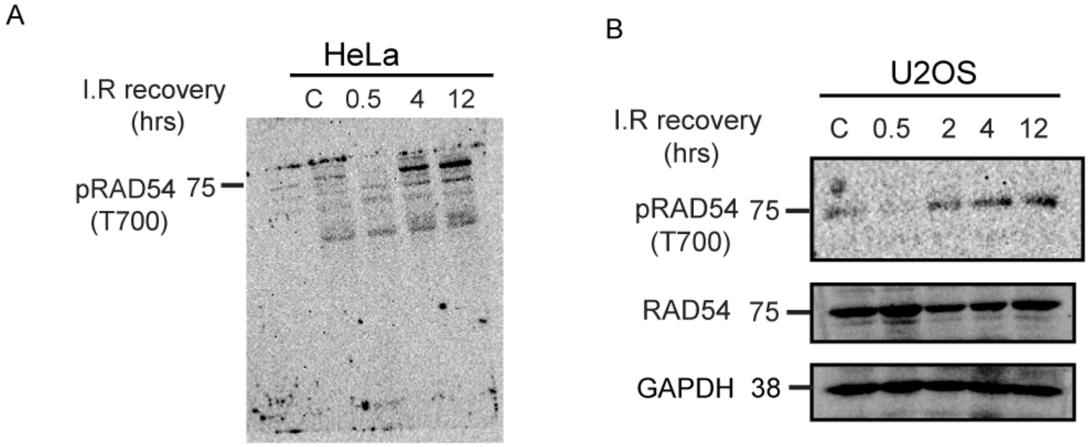


**Figure S10**: Phosphorylation of RAD54 increases post IR damage. A) Whole blot from HeLa cell lysate during increasing IR recovery probed with pRAD54 (T700) antiserum. B) U2OS cells treated with IR and allowed to recover as indicated. Western blot probed with pRAD54 (T700), RAD54 and GAPDH (loading control).

Supplementary Figure S11

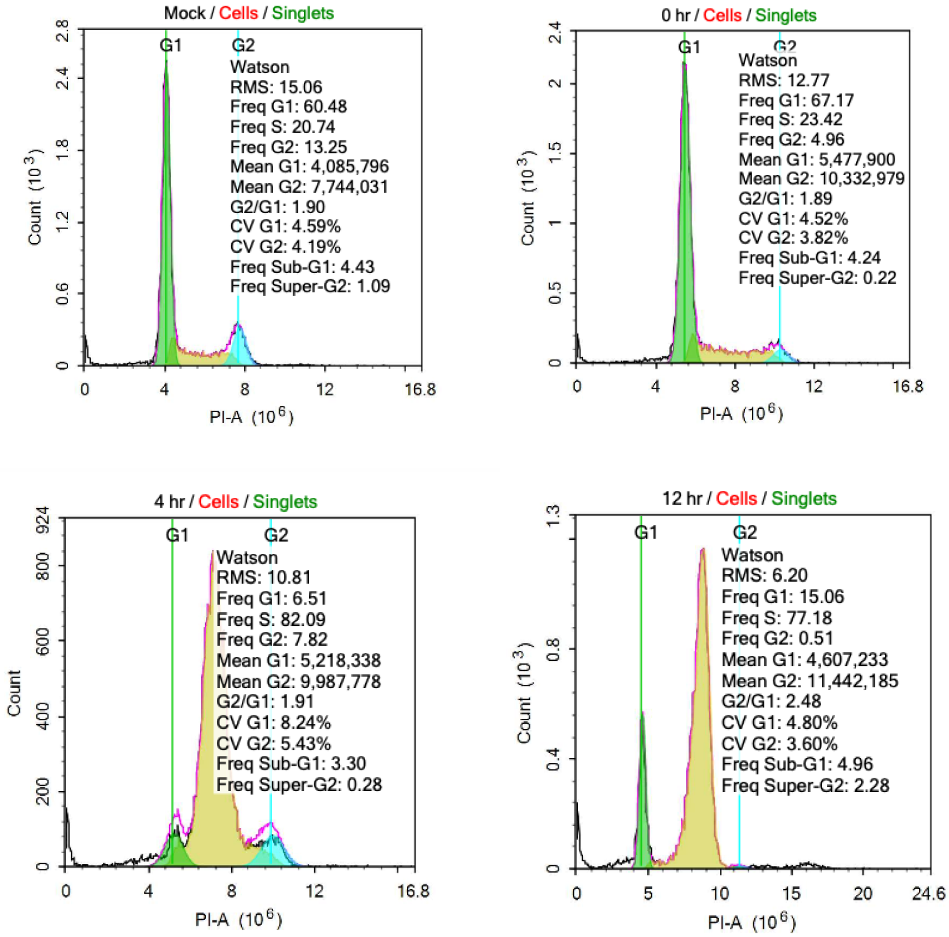


**Figure S11:** Cell cycle analysis of asynchronous HeLa cells (top left) and synchronized HeLa cells in S-phase and released for 0, 4 and 12hrs.

Supplementary Figure S12

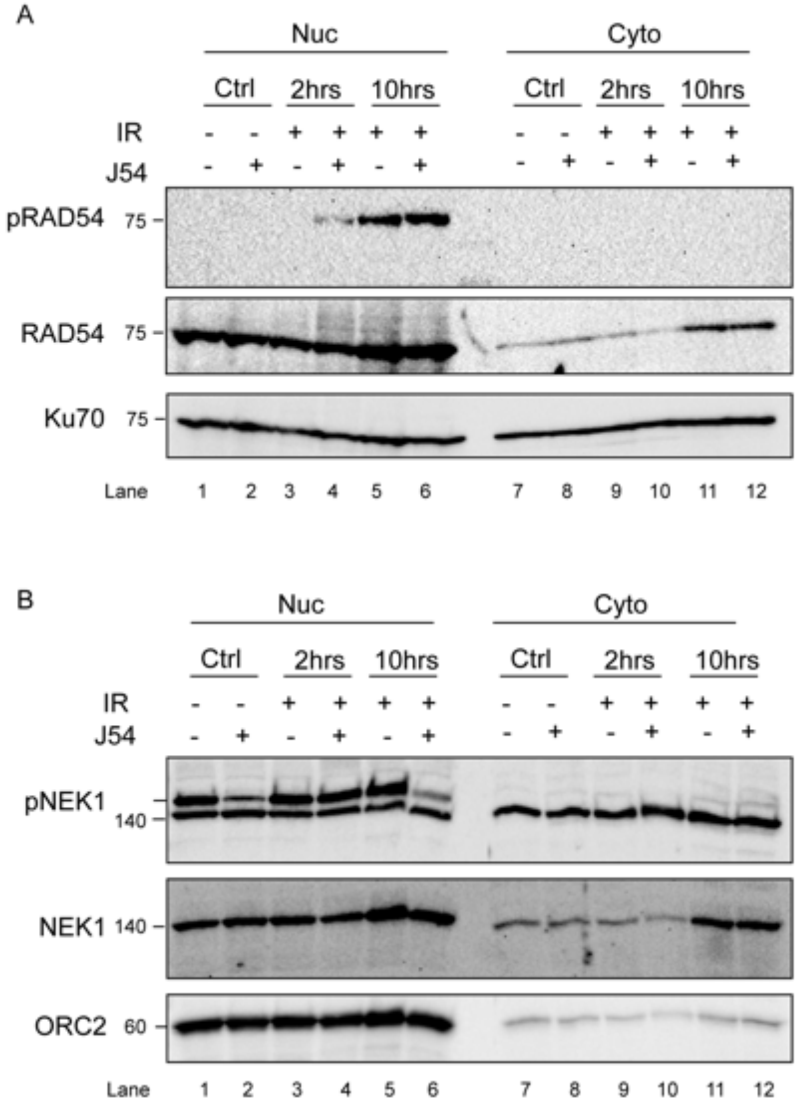


**Figure S12**: Phosphorylated RAD54 is nuclear post-irradiation. A) pRAD54 (T700) antibody used to probe nuclear and cytoplasmic fractions of cells treated without or with J54 induced with IR and allowed to recover for indicated times. Total RAD54 probed and Ku70 as loading control. Note that Ku70 is both nuclear and cytoplasmic. B) pNEK1 (T141) was probed in same experimental conditions as above (A). Total NEK1 was probed to show the preferential localization of the protein in nucleus. ORC2 was probed for chromatin loading control in same condition.

Supplementary Figure S13

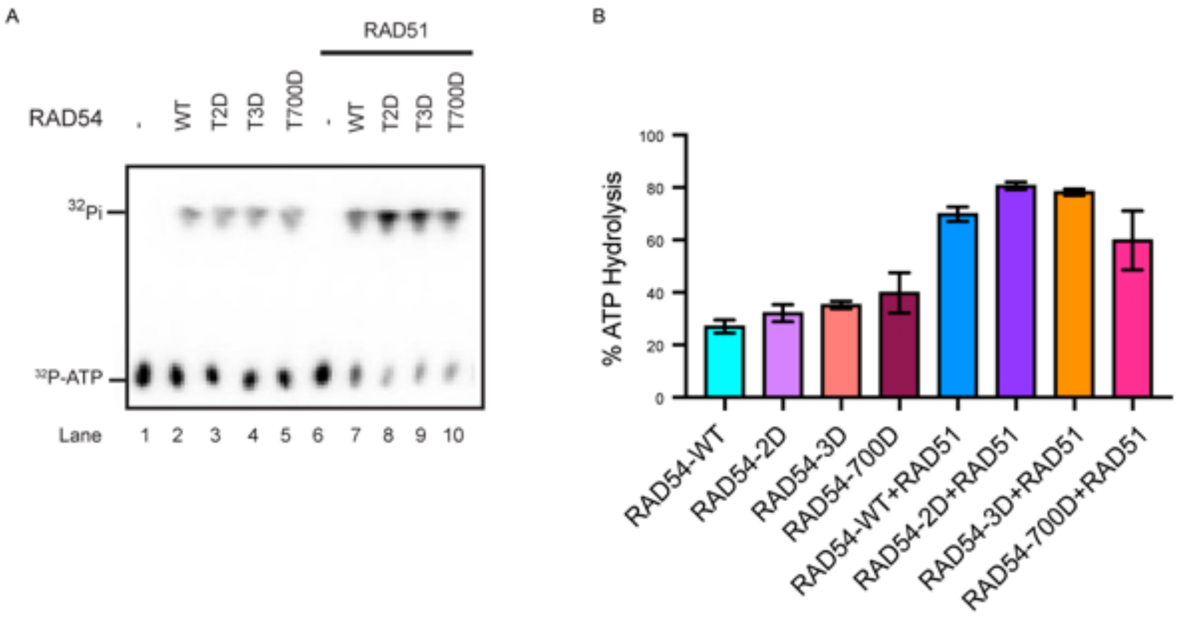


**Figure S13**: ATPase activity of RAD54 mutants does not alter at low salt concentration (22.5mM KCl). A) ATP hydrolysis activity of RAD54 without RAD51 (lane 2-5) and with RAD51 (lane 7-10). B) Quantification shown for assay in (A).

Supplementary Figure S14

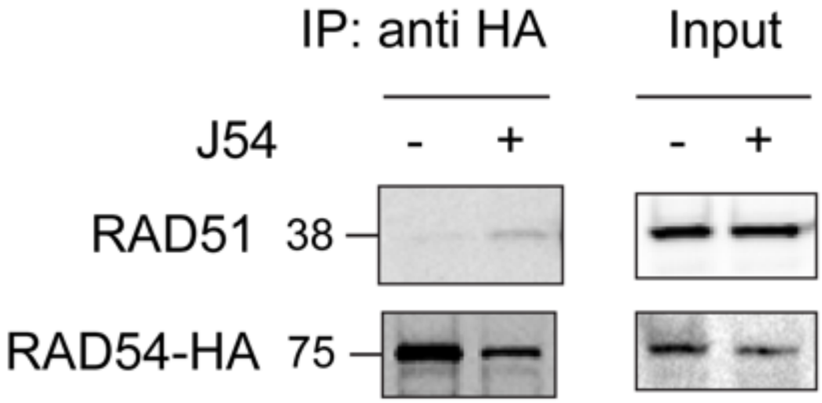


**Figure S14**: RAD51 interacts with RAD54 when TLK1 is inhibited (with J54, 10μM). Western blot to show the interaction between RAD54-HA and RAD51 in presence or absence of J54, without IR. RAD54-HA precipitated by anti-HA agarose beads. Input lanes show 5% of total protein loaded for IP.

Supplementary Figure S15

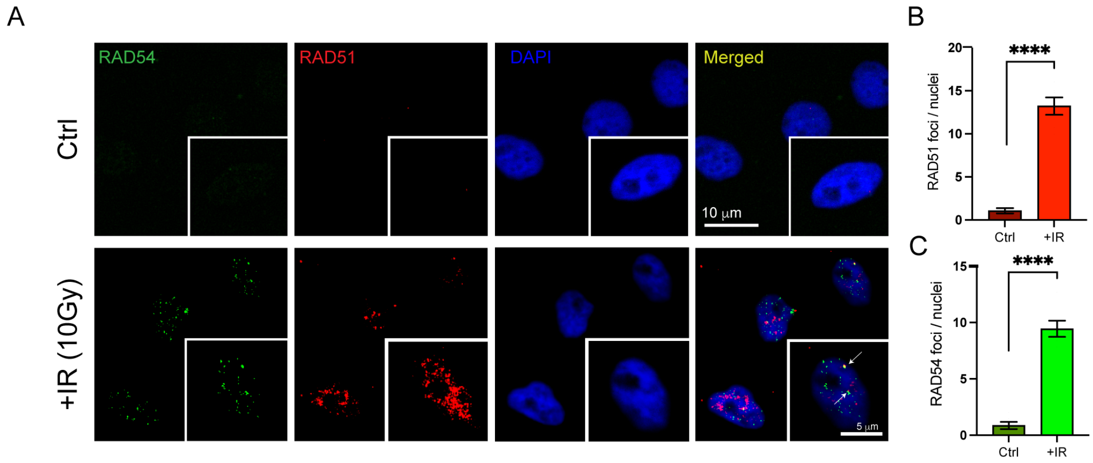


**Figure S15**: RAD54 and RAD51 forms discrete foci following IR. A) HeLa cells treated with IR (10Gy) and allowed to recover for 4hrs. RAD54 (green) and RAD51 (red) co-stained with antibody. Nucleus stained with DAPI. Scale bar is 10μm (inset 5μm). White arrows indicate yellow colocalization dots. B) Quantification shown for RAD51 foci per nuclei. C) Quantification shown for RAD54 foci per nuclei.

Supplementary Figure S16

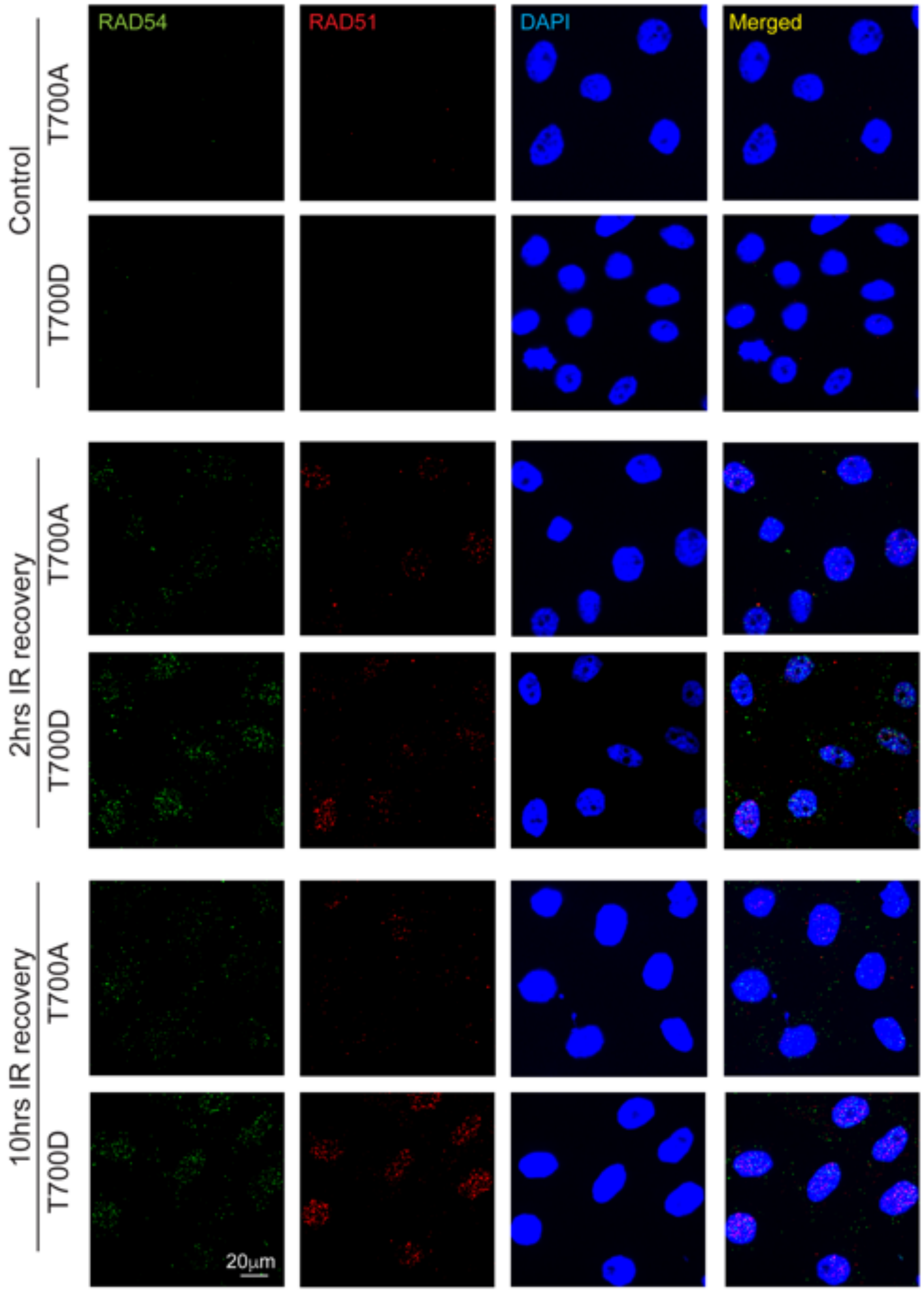


**Figure S16**: RAD54 and RAD51 co-stained in RAD54-T700A and RAD54-T700D cells without DNA damage (Control) and after indicated time of recovery post IR (10Gy) induction. T700D foci count is higher than T700A post irradiation at 10hrs post-IR which indicates RAD54-T700D foci dissolution is delayed compared to T700A.

Supplementary Figure S17

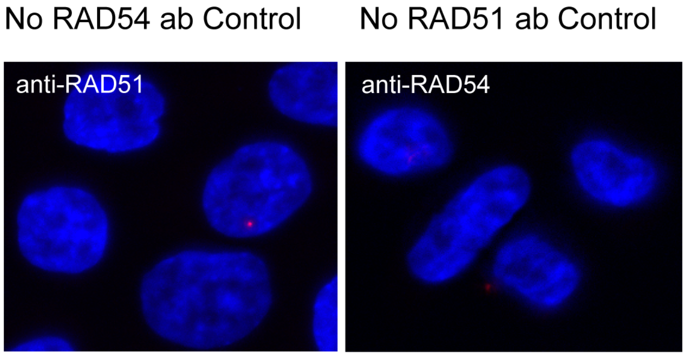


**Figure S17**: PLA assay used in samples where either of the RAD54 or RAD51 antibodies (negative control) were absent.

Supplementary Figure S18

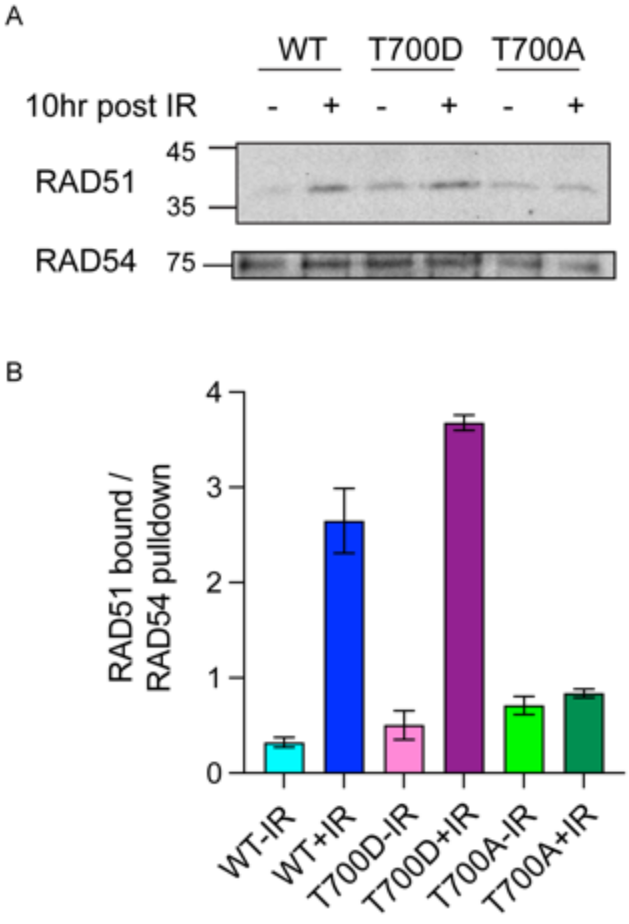


**Figure S18**: Phosphomutants of RAD54-T700 and RAD51 association increases post IR induction at 10hrs post IR. A) RAD54-T700D or T700A and WT cells were irradiated and recovered till 10hrs. Immunoprecipitation of RAD54-HA using HA-antibody was done and immunoblot probed for RAD51 and RAD54. B) Quantification of RAD51 bound to RAD54 fraction is plotted in graph.

Supplementary Figure S19

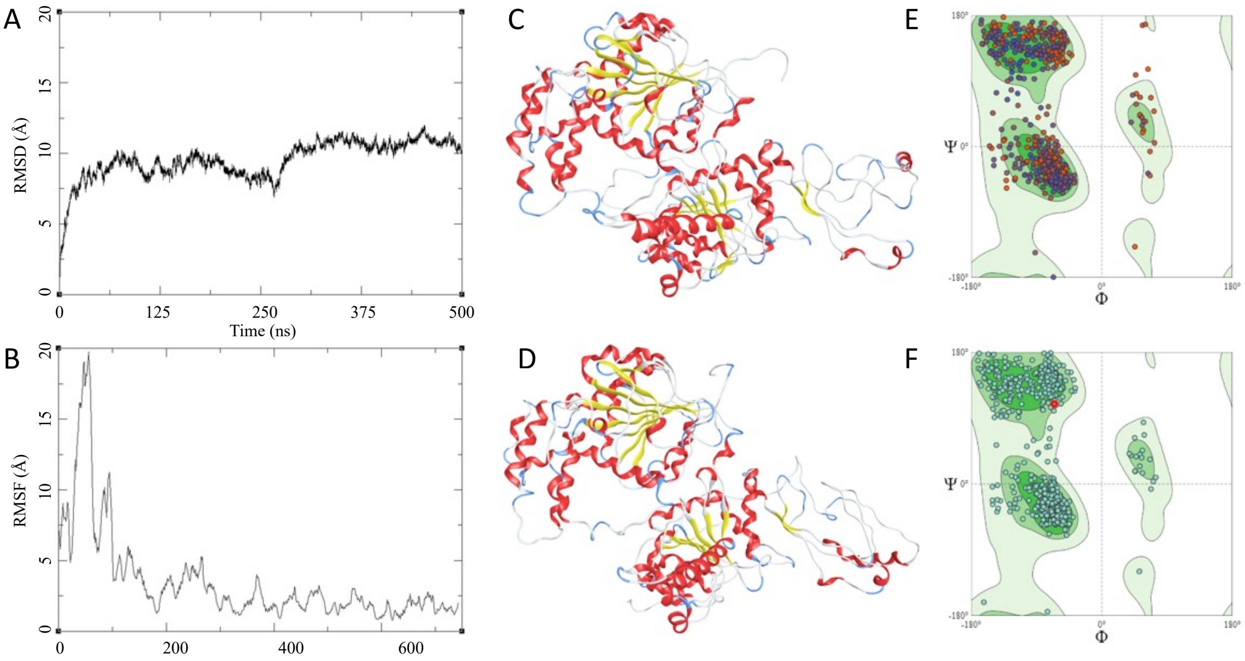


**Figure S19**: RAD54-WT: (a) RMSD of the RAD54-WT over 500 ns; (b) RMSF of the RAD54-WT over 500 ns; (c) Initial state of RAD54-WT at the beginning of 500 ns; (d) Final state of RAD54-WT at the end of 500 ns; (e) Ramachandran plot for the initial state of the RAD54-WT (MolProbity Score1.72, Qmean= -2.98); (f) Ramachandran plot for the final state of the RAD54-WT (MolProbity Score1.55, Qmean= -2.52).

Supplementary Figure S20

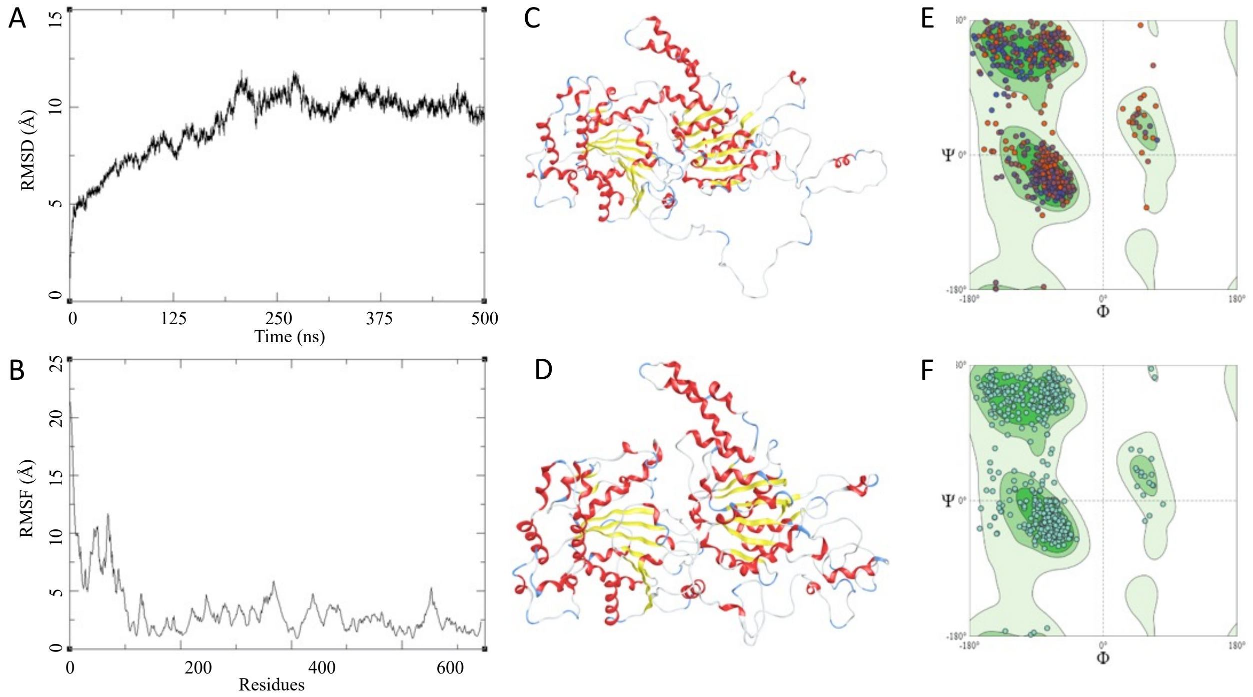


**Figure S20**: RAD54-T700D: (a) RMSD of the RAD54-T700D over 500 ns; (b) RMSF of the RAD54-T700D over 500 ns; (c) Initial state of RAD54-T700D at the beginning of 500 ns; (d) Final state of RAD54-T700D at the end of 500 ns; (e) Ramachandran plot for the initial state of the RAD54-T700D (MolProbity Score1.45, Qmean= -2.90); (f) Ramachandran plot for the final state of the RAD54-T700D (MolProbity Score1.56, Qmean= -2.47).

Supplementary Figure S21

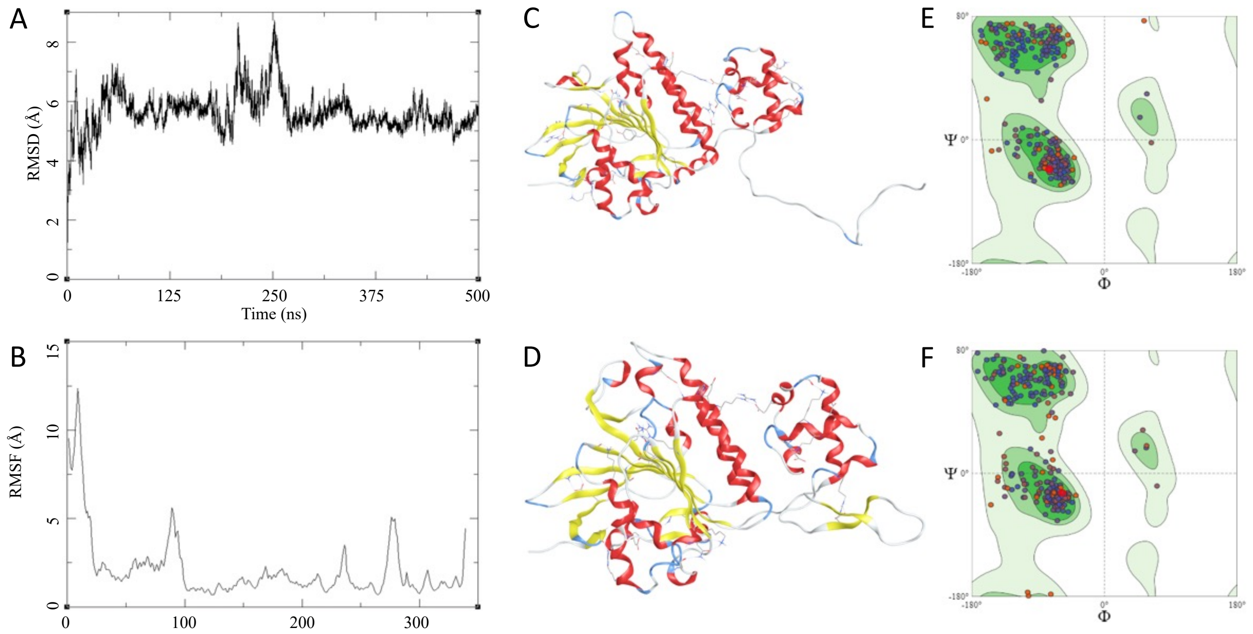


**Figure S21**: RAD51, (a) RMSD of the RAD51 over 500 ns; (b) RMSF of the RAD51 over 500 ns; (c) Initial state of RAD51 at the beginning of 500 ns; (d) Final state of RAD51 at the end of 500 ns; (e) Ramachandran plot for the initial state of the RAD51 (MolProbity Score1.41, Qmean= -1.59); (f) Ramachandran plot for the final state of the RAD51 (MolProbity Score1.27, Qmean= -1.54).
